# Vaccine genetics of IGHV1-2 VRC01-class broadly neutralizing antibody precursor naïve human B cells

**DOI:** 10.1101/2021.03.01.433480

**Authors:** Jeong Hyun Lee, Laura Toy, Justin T. Kos, Yana Safonova, William R. Schief, Corey T. Watson, Colin Havenar-Daughton, Shane Crotty

## Abstract

A successful HIV vaccine must overcome the hurdle of being able to activate naïve precursor B cells encoding features within their germline B cell receptors (BCR) that allow recognition of broadly neutralizing epitopes. Knowledge of whether broadly neutralizing antibody (bnAb) precursor B cells are circulating at sufficient frequencies within individuals in communities heavily impacted by HIV may be important. Using a germline-targeting eOD-GT8 immunogen and high-throughput droplet-based single cell BCR sequencing, we demonstrate that large numbers of paired BCR sequences from multiple donors can be efficiently screened to elucidate precursor frequencies of rare, naïve VRC01-class B cells. The results indicate that IGHV1-2 alleles incompatible with VRC01-class responses are relatively common in various human populations, and germline variation within IGHV1-2 associates with gene usage frequencies in the naïve BCR repertoire.

## INTRODUCTION

Broadly neutralizing antibodies (bnAbs) are present in a minority of patients chronically infected with human immunodeficiency virus-1 (HIV)^1^. These antibodies achieve neutralization breadth and potency against diverse circulating clinical strains by accruing high numbers of somatic hypermutations (SHM), allowing B cells to efficiently bind to conserved epitopes on the HIV Envelope viral spike protein (Env). BnAb structural and genetic analyses have shown that many bnAb features required for broad and potent neutralization, such as specific CDR lengths^2–5^ and certain amino acid residues at fixed positions defined by immunoglobulin (IG) variable (V), diversity (D), or joining (J) gene usage^6,7^, are predetermined by recombined naïve B cell receptors (BCRs). The majority of B cells in the human repertoire do not have BCRs with the potential to become HIV bnAbs. Thus, vaccine priming of rare bnAb precursor B cells likely require custom immunogens designed to bind specifically to targeted precursors. Making the problem even more challenging, inferred germline (iGL) precursors for many potent HIV bnAbs, have been found to have very low or no detectable affinity for wild-type HIV Env^8–14^, and wildtype Env immunogens have not succeeded in eliciting bnAb responses^15^. This lack of affinity of bnAb precursors for wildtype HIV Env remains one of the main impediments in neutralizing antibody directed HIV vaccine efforts.

One theoretical approach to recapitulating bnAb responses via vaccination involves priming with an immunogen that has exceptionally high affinity for bnAb precursors, then sequentially introducing more native Env-like immunogens to drive bnAb class SHMs^16^. Such priming immunogens are fittingly described as GT priming immunogens^17^, and a sequential vaccination strategy anchored by these priming immunogens has been described as “germline-targeting vaccine design”^18,19^. Several GT priming-immunogens have been designed specifically to bind the inferred germline (iGL) versions of known bnAbs with high affinity^12,13,18–23^. For the GT priming immunogens to be efficacious, at least two biological prerequisites must be met; the majority of the human population must have the genetic capacity to encode the targeted germline B cells^13,20,24,25^, and the frequency of such B cells needs to be high enough that they can respond to the immunogen while simultaneously competing against off-target B cells during maturation^13,15,19,20,26–28^.

Using carefully controlled mouse models, it has been shown that parameters that can be used to predict how well an immunogen will perform include: the target B cell precursor frequency, the monovalent affinity of the precursor B cell to the immunogen, and the avidity/multivalency of the immunogen^24,26,29–31^. Because the starting precursor frequency of target B cells in humans cannot be manipulated, it is a key parameter according to which an immunogen need be iteratively designed in order to increase the target affinity. We have previously developed a strategy to directly quantify bnAb B cell precursor frequencies from the human B cell repertoire, by using high affinity GT-probes to isolate antigen-specific naïve B cells from blood of healthy individuals^19,20,32^. One class of bnAbs that were analyzed by this method was precursors to Env CD4 binding site (CD4bs) targeting bnAbs, termed VRC01-class^33^. VRC01-class BCRs are identifiable by the use of alleles at the IG heavy chain (IGH) variable gene IGHV1 −2 paired with a light chain (LC) with a short complementarity determining region (CDR) 3 of 5-amino acids (AA)^34–36^. The engineered outer domain (eOD) derived GT immunogen eOD-GT8 is designed to VRC01 -class B cells. eOD-GT8 was able to bind VRC01 -class precursor naive B cells in human blood samples^20,32^, and eOD-GT8 was able to activate germline naïve VRC01 -class B cells at 1 in 1 million precursor frequency in a small animal model^26^. These were among the key findings that helped advance eOD-GT8 60mer to phase 1 clinical trial as a GT HIV vaccine candidate prime (NCT03547245).

In previous human B cell repertoire screening for antigen specific naïve B cells, antigen-probe binding B cells were single cell sorted, then subjected to nested polymerase chain reactions (PCR) performed separately for the BCR HC (IGHV), and LC IG kappa (IGKV) and lambda (IGLV) genes. While this method can be efficient when querying a small number of B cells, it becomes overly time consuming and costly as the number of analyzed BCR sequences increases. With the improved droplet single cell RNA sequencing (scRNA-seq) technologies, it is possible to efficiently recover single cell transcriptomic data from bulk sorted cells. Recently, several groups have performed high-throughput antigenspecific B cell repertoire sequencing using the single cell immune profiling platform from 10x Genomics^37–40^. However, no studies to our knowledge have used this technology to specifically sort rare antigen-specific naïve human B cells. Here, we used droplet-based scRNA-seq to obtain HC and LC paired VRC01-class naïve human BCR sequences and demonstrated that this method can reliably identify rare antigen-specific B cells. Additionally, we used these data along with other samples from ethnically diverse population cohorts to analyze the human population genetics of our VRC01 – class bnAb targeting vaccine.

## RESULTS

### Identification of naïve VRC01-class B cells using tetramer probes and high-throughput sequencing

Several CD4bs GT immunogens have been designed to bind VRC01-class precursor BCRs, some of which use the engineered outer domain of gp120 (eOD) as the base molecule^13,20^. Of these, eOD-GT8 has an average of ~6 nM affinity to several iGL VRC01-class bnAbs^20^. By tethering biotinylated eOD-GT8 monomers to fluorescently labeled streptavidin (SA) to generate tetramers, we previously isolated CD4bs-specific B cells from the human naïve B cell repertoire, and identified VRC01-class naïve B cells by single cell BCR sequencing to reveal that these cells are found in healthy humans at a frequency of ~1 in 300,000 naïve B cells^20,32^.

To determine whether droplet-based scRNA-seq was applicable to sequencing rare antigen-specific B cells, we sorted eOD-GT8-specific naïve B cells from PMBCs of three independent healthy donors and used the 10X Genomics Chromium platform to obtain BCR sequences (Fig. 1). As in our previous experiments, antigen-specific cells were defined as those that bound eOD-GT8:SA on two different fluorescent probes while not binding the eOD-GT8 knockout- II (eOD-GT8^KO^) probe (Fig. 1 a-c), which is identical to eOD-GT8 except for mutations in the CD4bs that prevents VRC01 – class B cells and their respective iGL B cells from binding^29,32^. From here on, eOD-GT8^+^ refers to cells that bind the eOD-GT8 probe on two different fluorophores.

**Figure 1.**
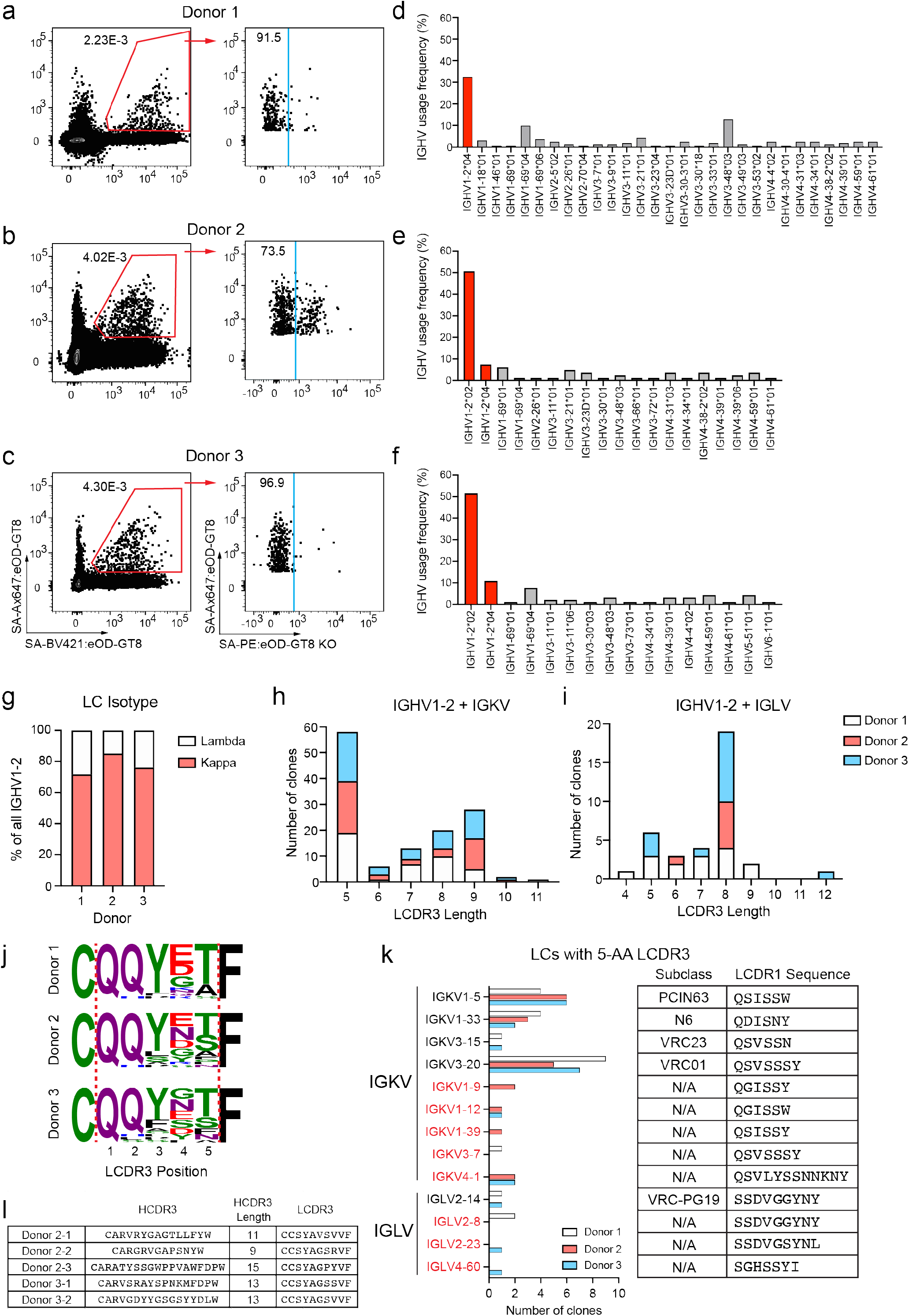
eOD-GT8 tetramer sorted VRC01-class naive B cells. B cells enriched from healthy donor PBMCs were stained with biotinylated eOD-GT8 probes formed into tetramers by binding to SA, and sorted for eOD-GT8^+^eOD-GT8^KOneg^ naïve B cells, followed by BCR sequencing. **(a-c)** Flow cytometry of eOD-GT8^+^ eOD-GT8^KOneg^ cells in donor 1 **(a)**, donor 2 **(b)**, and donor 3 **(c)**. Cells in the blue gate were sorted. **(d-f)** IGHV gene usage distribution of paired eOD-GT8^+^eOD-GT8^+^/eOD-GT8^KOneg^ naïve B cells from donor 1 **(d)**, donor 2 **(e)**, and donor 3 **(f)**. Bars corresponding to VH1-2 genes are colored in red. **(g)** Isotype distribution of LCs coexpressed with IGHV1 −2. **(h)** LCDR3 lengths of IGKV LCs coexpressed with IGHV1 −2. **(i)** As in (h), but for IGLV LCs. **(j)** LCDR3 sequences of 5-AA IGKV LCs. **(k)** IGKV and IGLV usage distribution among eOD-GT8^+^, 5-AA LCDR3 VRC01-class naïve B cells and their germline encoded LCDR1 sequences. IGKV and IGLV genes shown in black are genes observed in known VRC01-class bnAbs, whereas those indicated in red have not been observed. **(l)** IOMA-class B cells identified using tetramer probes.

Droplet scRNA-seq functions by capturing a single cell along with a uniquely barcoded bead within a droplet. Occasionally more than one cell is captured per partition. Thus, we first cleaned up our annotated paired VDJ sequences. Cell barcodes associated with doublets were identified by the presence of two HC contigs, and/or two LC contigs of the same isotype (e.g. two IGLV or two IGKV contigs). Cells that did not have a HC-LC pair were also eliminated from analysis. Because the primers in this system were also designed to engage all IGH constant region genes, we were able to identify IgA^+^ or IgG-subclass^+^ BCRs that escaped the dump gate during sorting and remove them from analysis. Lastly, cells sorted from donors 2 and 3 were multiplexed with other sort samples by hashtag feature barcoding^41^, and cell barcodes associated with dual high-hashtag counts were removed. The final number of recovered paired BCR sequences were 163, 81, and 114 in donors 1 through 3 respectively (Table 1). The IGHV1-2 gene was highly enriched among paired BCR sequences. Across the three donors, between 32-60% of the HC were IGHV1-2 (Fig. 1d-f). 71-86% of LC paired with IGHV1-2 were IGKV (Fig. 1g). 37-43% of the LCs (42-50% IGKV and 0-21% IGLV) paired with a IGHV1-2 HC harbored a short 5-AA CDR3 (Fig. 1h-i). These frequencies of eOD-GT8 tetramer^+^ total IGHV1-2 HCs, and the proportion of 5-AA LCDR3 LCs among IGHV1 −2 HCs, were comparable to what was observed in a Sanger sequencing-based study (67% and 33% respectively)^32^. The precursor frequency of VRC01-class naïve B cells, defined as the proportion of IGHV1-2 HC paired with a LC with a 5-AA LCDR3 among total naïve B cells, ranged between 1 in 0.13 million to 0.25 million B cells (Table 1), similar to the previously reported frequency of 1 in 0.3 million B cells^32^. Most IGKV VRC01 -class precursors had the VRC01 -class bnAb signature LCDR3 sequence of QQYXX and E/N/Q at position 4 (Fig. 1j). The sequences of VRC01 -class naïve B cells isolated from donors 1 −3 are provided in Supplementary Table 1.

**Table 1.**
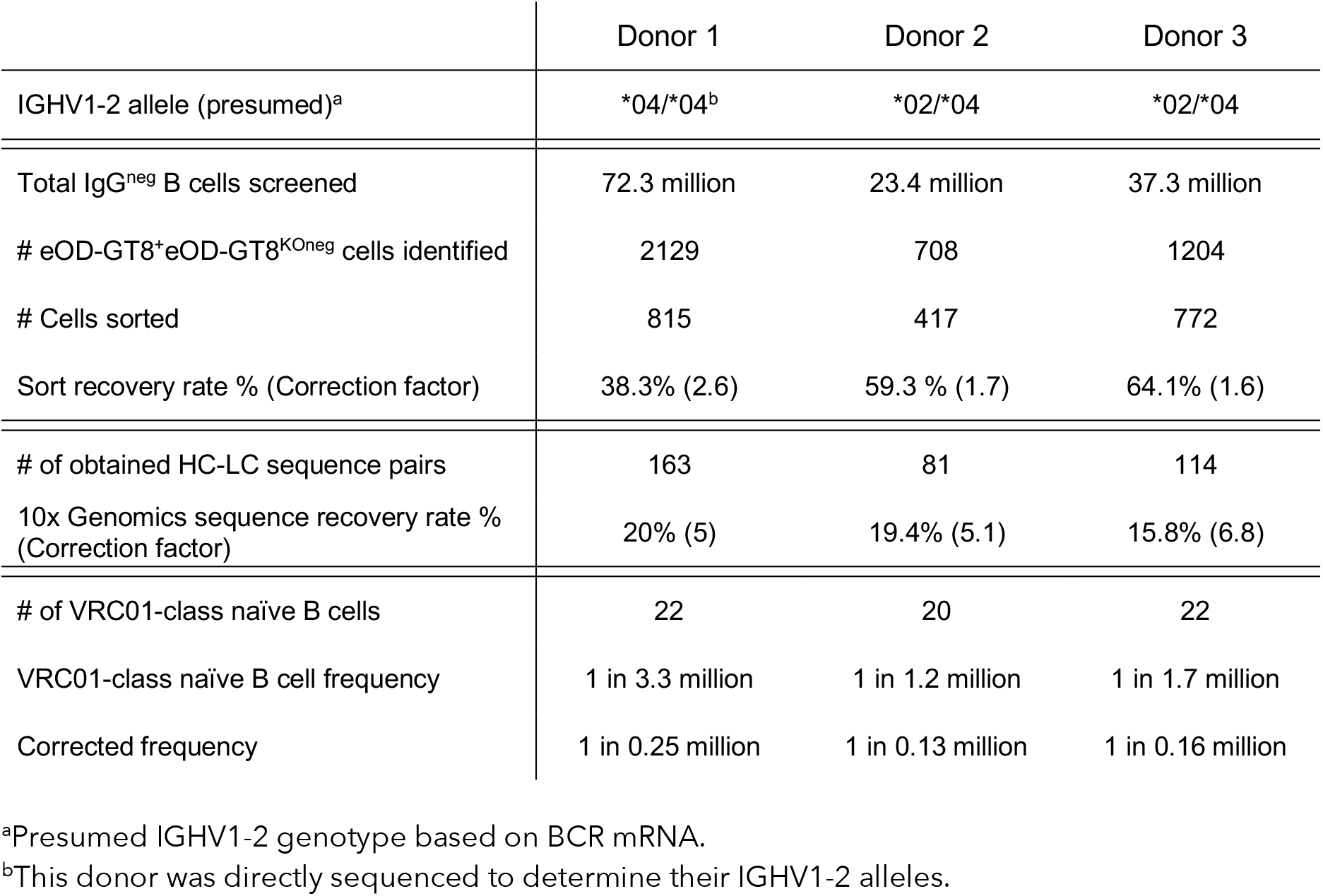
VRC01-class naïve B cell precursor frequency calculated for cells sorted with eOD-GT8 tetramers

### Diverse range of VRC01-class precursor LCs

The LC variable gene usage among known VRC01 -class antibodies allows room for diversity, but it is desirable for the germline encoded LCDR1 to be short in length or rich in flexible glycine or serine residues, as this region of the antibody is critical in accommodating the conserved N276 N-linked glycan near the CD4bs^35^. The majority of VRC01 – class bnAbs utilize IGKV, possibly because the average length of CDRL1 among IGKV genes (6-AA) of circulating naïve B cells in the human repertoire are shorter than IGLV genes (9-AA)^32^. In sequences from 3 donors, 70-95% of VRC01 – class BCRs possessed IGKV genes expressed by known VRC01-class bnAb subtypes (VRC01-subclass: IGKV3-2 0^35^; PCIN64-subclass: IGKV1-5^4^; N6-subclass: IGKV1-33^42^; VRC23-subclass: IGKV3-15^43^). BCRs expressing IGKV1-12, IGKV1-39, or IGKV3-7 were observed for the first time. Almost all of the IGKV sequences not observed among known VRC01 – class bnAbs had glycine and serine containing 6-7 AA LCDR1s (Fig. 1k).

Of the few IGLV VRC01-class naïve B cell sequences, the average LCDR1 length was 9-AA, in accordance with the average LCDR1 length of human IGLV genes. Only one clone expressed IGLV4-60, which has a short LCDR1 length of 6-AA, but all of the IGLV LCDR1 sequences were enriched with glycine and serine residues (Fig. 1k). Notably, we identified two IGLV2-14 clones from two independent donors, this being the first instance of having isolated a VRC01-class precursor naïve B cell belonging to the VRC-PG19 bnAb subclass^7,32^.

Previously it was shown that IOMA-class naïve B cells could be isolated using eOD-GT8 tetramer probes^32^. IOMA is a CD4bs bnAb that has a IGHV1 −2 HC but utilizes a IGLV2-23 LC with an 8-AA LCDR3 and a slightly different mode of binding compared to classic VRC01-class bnAbs^44^. In our current study, 5 IOMA-class B cells were obtained from two donors (Fig. 1l). Overall, these results demonstrate that bulk-sorting followed by droplet scRNA-seq can be a productive approach to identify BCRs of vaccine-specific naïve B cells.

### High-avidity antigen probes increase capture of off-target B cells

Binding of low affinity B cells to antigens can be dramatically augmented by using multimeric proteins to improve avidity^13,26,45–47^. For example, high avidity eOD-GT8 60mer nanoparticle fluorescent probes could identify an increased frequency of eOD-GT8^+^ B cells by flow cytometry, of which approximately 90% did not co-stain for eOD-GT8^KO^ 60mer^32^, potentially selecting for lower affinity VRC01-class naïve B cells that could not be identified using tetramer probes. However, eOD-GT8 60mer^+^ binding B cells were not previously BCR sequenced. Thus, the efficiency in isolating precursor B cells by high-avidity nanoparticle probes is unknown. To probe the eOD-GT8 60mer^+^ naïve B cell repertoire, we performed BCR sequencing of cells from three healthy donors sorted using eOD-GT8 60mer and eOD-GT8^KO^ 60mer probes (Fig. 2).

**Figure 2.**
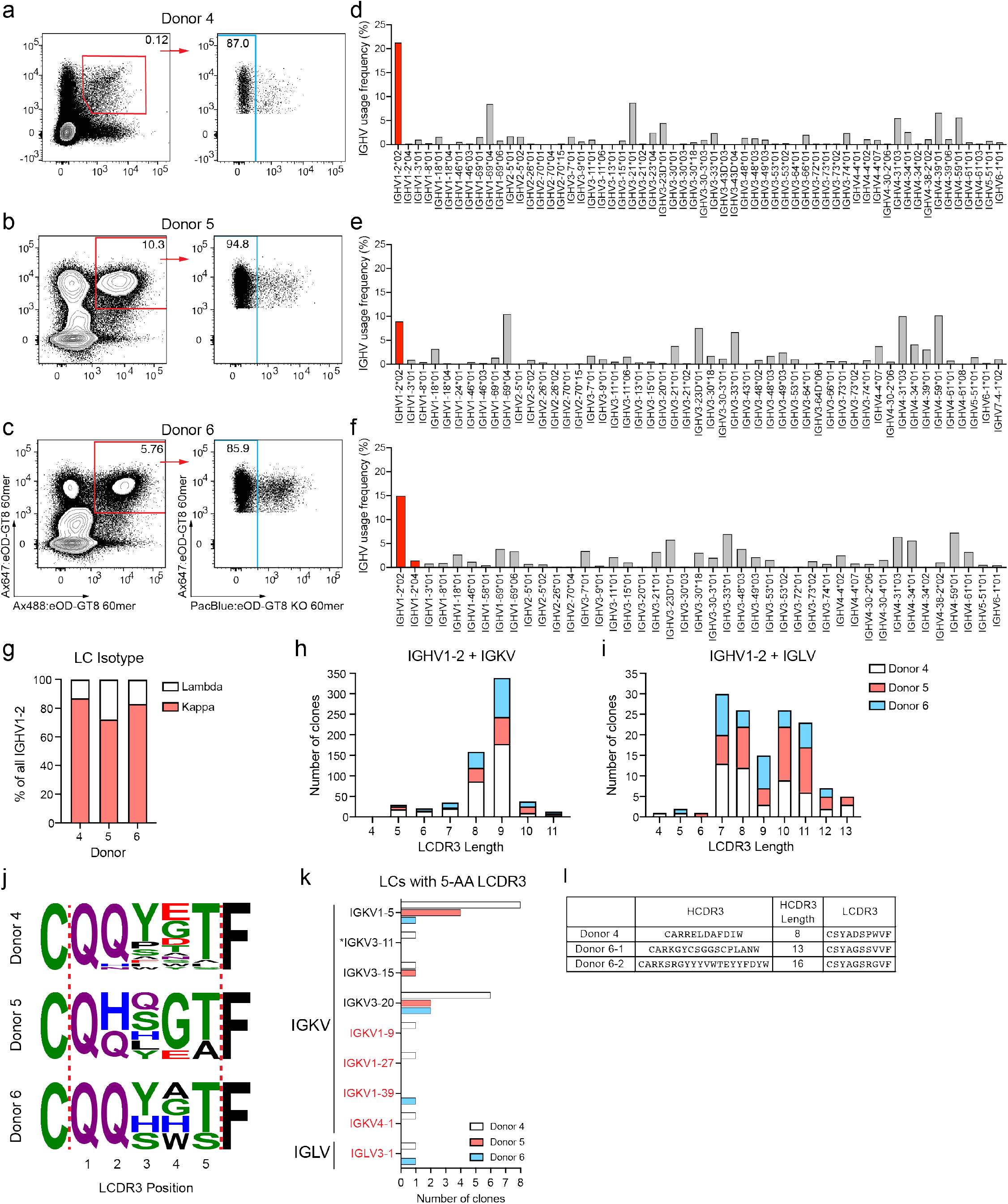
eOD-GT8 60mer sorted VRC01-class naive B cells. B cells enriched from healthy donor PBMCs were stained with eOD-GT8 60mer probes directly conjugated with fluorophores and sorted for eOD-GT8 60mer^+^eOD-GT8 60mer^KOneg^ naïve B cells, followed by BCR sequencing. **(a-c)** Flow cytometry of eOD-GT8 60mer^+^eOD-GT8 60mer^KOneg^ cells in donor 4 **(a)**, donor 5 **(b)**, and donor 6 **(c)**. Cells in the blue gate were sorted. **(d-f)** IGHV gene usage distribution of paired eOD-GT8 60mer^+^eOD-GT8 60mer^KOneg^ naïve B cells from donor 4 **(d)**, donor 5 **(e)**, and donor 6 **(f)**. Bars corresponding to IGHV1-2 genes are colored in red. **(g)** Isotype distribution of LCs coexpressed with IGHV1 −2. **(h)** LCDR3 lengths of IGKV LCs coexpressed with IGHV1 −2. **(i)** As in (h), but for IGLV LCs. **(j)** LCDR3 sequences of 5-AA IGKV LCs. **(k)** IGKV and IGLV usage distribution among eOD-GT8^+^, 5-AA LCDR3 VRC01-class naïve B cells. Color coding is as in (Fig. 1k). IGKV3-11 indicated by an asterisk has not been directly observed in VRC01-class bnAbs, although the LC of VRC01 was originally annotated as IGKV3-11^33^ as it is extremely similar to IGKV3-20. **(l)** IOMA-class naïve B cells sorted with eOD-GT8 60mer probes.

Cells were stained in two different ways. PBMCs from donor 4 were first enriched for B cells, then stained with fluorescent probes and antibodies as was done for all previous tetramer probe experiments (Fig. 2a). For donors 5 and 6, total PBMCs were instead incubated with AlexaFluor647-conjugated eOD-GT8 60mer first, then enriched for AlexaFluor647^+^ cells followed by staining with Alexafluor488-eOD-GT8 60mer, Pacific-Blue-eOD-GT8^KO^ 60mer, and antibodies (Fig. 2b, c). By doing so, pre-sort eOD-GT8 60mer^+^ cells were enriched by ~40 fold (Fig. 2a-c, Supplementary Fig. 1 a). Regardless of the sample preparation method used, a substantially larger fraction of naïve B cells stained eOD-GT8 60mer^+^eOD-GT8^KO^ 60mer^neg^ than when tetramers were used. As a result, a much higher total number of naïve B cells could be sorted per donor (Table 2). More than 1000 paired BCR sequences were obtained from each donor, even though the sequence recovery rates were slightly reduced relative to the input number of cells compared to tetramer sorts (Table 2).

**Table 2.** VRC01-class naïve B cell precursor frequency calculated for cells sorted with eOD-GT8 60mers.

|  | Donor 4 | Donor 5 | Donor 6 |
| --- | --- | --- | --- |
| IGHV1-2 allele (presumed)<sup>a</sup> | *02/*04 | *02/*02 | *02/*04 |
| Total IgG<sup>neg</sup> B cells screened | 25.6 million | 246,890 | 419,167 |
| Total B cells corrected for Ax647 enrichment<sup>b</sup> | N/A | 9.9 million | 16.8 million |
| # eOD-GT8<sup>+</sup>eOD-GT8<sup>KOneg</sup> cells identified | 27,902 | 24,054 | 20,737 |
| # Cells sorted | 16,076 | 11,132 | 8,727 |
| Sort recovery rate % (Correction factor) | 57.6% (1.7) | 46.3 % (2.2) | 42.1% (2.4) |
| # of obtained HC-LC sequence pairs | 1,786 | 2,039 | 1,249 |
| 10x Genomics sequence recovery rate % (Correction factor) | 11.1 % (9) | 18.3% (5.5) | 14.3 % (7.0) |
| # of VRC01-class naïve B cells | 20 | 7 | 5 |
| VRC01-class naïve B cell frequency | 1 in 1.28 million | 1 in 1.42 million | 1 in 3.35 million |
| Corrected frequency | 1 in 0.08 million | 1 in 0.12 million | 1 in 0.20 million |

<sup>a</sup>Presumed IGHV1-2 genotype based on BCR mRNA. Donor 5 may have a second allele other than \*02 not identified by the sorted B cells.
<sup>b</sup>B cells were not stained prior to AlexaFluor647 enrichment. Therefore, we cannot back-calculate the true total number of IgG<sup>neg</sup> B cells in donor 5 and donor 6 samples. Since the majority of cells bound to eOD-GT8 60mer probes were not VRC01-class antibodies, we estimated the normal proportion of B cells that are eOD-GT8 60mer<sup>+</sup> by averaging frequencies from 60mer sorted samples; from 10x Genomics donor 4, the IGHV1-2\*05 allele donor 8, and single cell sequencing donor 7 (Fig. 2a, Fig. 3b, Supplementary Fig. 1a, respectively). Frequency of B cells that are eOD-GT8 60mer<sup>+</sup>eOD-GT8 60mer<sup>KOneg</sup> post-AlexaFluor647 enrichment was estimated by averaging values from 60mer donors 5 and 6. Post-enrichment frequency was divided by pre-enrichment frequency to yield 40.15-fold enrichment of B cells.

Relative IGHV1-2 gene usage in each donor ranged from 9-22%, compared to 32-60% when using tetramer probes (Fig. 1d-f, Fig 2d-f). Of the IGHV1-2^+^ B cells, the ratio of IGKV to IGLV were similar to tetramer-sorted BCRs (Fig. 2g), but only 4.7% of IGKV and 1.5% of IGLV BCRs possessed 5-AA LCDR3s regardless of the LC isotype (Fig. 2h, i). Single cell Sanger sequencing of eOD-GT8 60mer^+^ B cells in another donor (donor 7) also found low frequencies of IGHV1-2 HCs. Of 103 B cells sequenced, 13% expressed IGHV1-2, of which just one cell had a 5-AA LCDR3 (Supplementary Fig. 1 b-d). Thus, the reduction in the number of identified VRC01 -class naïve B cells was not an artifact associated with the sequencing method.

All observed VRC01 -class naïve B cell sequences among eOD-GT8 60mer-bound B cells had canonical features of VRC01-class bnAbs (Fig. 2j, k). Three IOMA-class naïve B cells were also found among B cells sorted from donors 4 and 6 (Fig. 2l). These data suggest that the eOD-GT8 60mer binds rare VRC01 -class naïve B cells as designed, but the high avidity of the antigen captures low affinity B cells much more so than tetramers. Consistent with this conclusion, the final calculated VRC01-class B cell precursor frequencies identified by eOD-GT8 60mer probes ranged between 1 in 0.08 to 0.2 million naïve B cells, a range similar to the precursor frequency determined using eOD-GT8 tetramer probes (Table 2)^20,32^. The sequences of VRC01 -class naïve B cells isolated from donors 4-6 are provided in Supplementary Table 2.

### IGHV1 −2*05 is unable to bind the HIV CD4bs in a VRC01 -like manner

During our study, we identified one donor (donor 8) who had approximately 10-fold lower eOD-GT8^+^eOD-GT8^KOneg^ naïve B cells identified using tetramers, compared to previous donors (Fig. 1a-c, Fig. 3a). We sequenced tetramer sorted eOD-GT8^+^eOD-GT8^KOneg^ B cells by droplet scRNA-seq and found that surprisingly, none of BCRs expressed the IGHV1 −2 gene (Fig. 3c). When eOD-GT8 60mer nanoparticles were used as probes to stain cells from that same donor, the frequency of eOD-GT8 60mer^+^eOD-GT8 60mer^KOneg^ naïve B cells were similar to what was observed in other donors from whom we were able to isolate VRC01 -class naïve B cells (Fig. 2a-c, Fig. 3b, Supplementary Fig. 1 a^32^). Nearly 3000 paired BCR sequences were obtained from the 60mer sorted B cells, but only three B cell clones expressed an IGHV1 −2 gene (Fig. 3c). None of the three IGHV1 −2 B cells coexpressed a LC with a 5-AA LCDR3 (Fig. 3d-e, left-tailed population proportion hypothesis test: P < 0.0001). Out of all sequences, irrespective of the HC or probes used, three clones among 60mer sorted B cells had a 5-AA LCDR3. The three IGHV1-2 clones were annotated as having an *05 allele, predicted to be unsuitable as a VRC01 -class precursor due to a missing germline encoded W50 residue that forms a conserved interaction with N280 of gp120 in all VRC01-class bnAbs^13,34,36^. These observations led to the hypothesis that this donor had IGHV1-2 alleles with significantly reduced potential to develop VRC01-class bnAbs. The IGHV1-2 genotype of donor 8 was subsequently confirmed to be IGHV1-2*05/*05 by targeted PCR, cloning and sequencing (Supplementary Fig. 2a), incompatible with eOD-GT8 binding due to the missing W50^13,34,36^.

**Figure 3.**
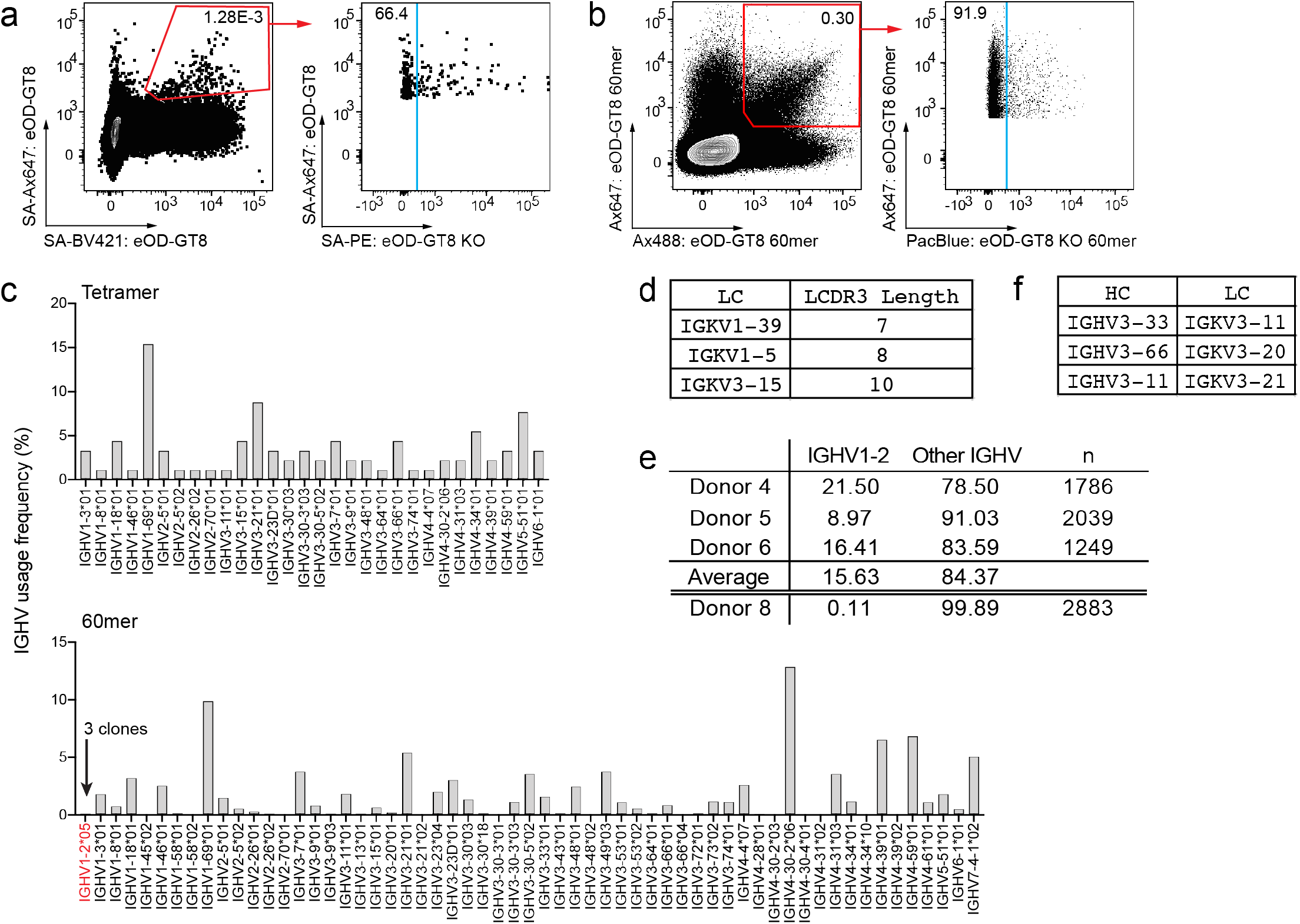
Some individuals do not have VRC01-class naïve B cells due to incompatible IGHV1-2 alleles. B cells were enriched from donor 8. PBMCs were stained for eOD-GT8 tetramer and eOD-GT8 60mer probe binding, and sorted for eOD-GT8^+^eOD-GT8^KOneg^ naïve B cells. **(a-b)** Flow cytometry of eOD-GT8^+^eOD-GT8^KOneg^ cells using tetramer probes **(a)**, or with **(b)** 60mer probes. Cells in the blue gate were sorted. **(c)** IGHV gene usage among paired BCR sequences derived from tetramer (upper, n=91) and 60mer (lower, n=2,856) sorted cells. **(d)** Characteristics of LCs paired to the three IGHV1 −2*05 clones detected from the 60mer sorted cells. **(e)** Frequency of IGHV1 −2 BCRs observed in eOD-GT8 60mer sorted datasets. **(f)** Three 5-AA LCDR3 harboring sequences were found among 60mer sorted cells. None of the paired HCs were IGHV1 – 2.

### Precursor frequencies of eOD-GT8 binding IGHV1-2 naïve B cells are affected by allelic variations

We noted that among our tetramer donors, eOD-GT8 tetramer donor 1 with the lowest frequency of IGHV1-2 B cells (32%) was homozygous for the IGHV1 −2*04 allele (Fig. 1d). In eOD-GT8 tetramer donors 2 and 3 who were both heterozygous for *02/*04, the majority of identified IGHV1-2 BCRs utilized the *02 allele (Fig. 1e-f). The VRC01-class naïve B cell precursor frequency in the homozygous IGHV1 −2*04 donor (tetramer donor 1; Supplementary Fig. 2b) was also lower by 2-fold, 1 in 0.25 million compared to an average of 1 in 0.145 million naïve B cells in donors 2 and 3 (Table 1). Of the 7 curated IGHV1 −2 alleles in IMGT, all currently known VRC01 -class bnAbs are thought to derive from IGHV1 – 2*02 allele^4,35,36^, even though the *03, *04 and *07 alleles have germline encoded W50, N58, and R71 residues (Kabat numbering) required for CD4bs recognition and thus have the potential to become VRC01 -class bnAbs^13,36^ (Fig. 4a). It is plausible that *01 and *03 alleles are old sequencing artifacts. The *07 allele has not been observed in donors so far, likely due to rarity, as the *07 allele was only recently annotated (GenBank: MN337615, unpublished). The *04 allele encodes a W66 in framework region (FWR) 3 in place of an arginine found in other IGHV1-2 alleles. Arginine is the preferred residue at position 66 among annotated productive human IGHV genes (Fig. 4b). The next most common variants in this position are Q66 and H66, which both retain polar side chains. The hydrophobic tryptophan residue exposed on the surface of the antibody may impact the solubility of the W66 harboring BCR, thereby affecting development of IGHV1 −2*04 B cells.

**Figure 4.**
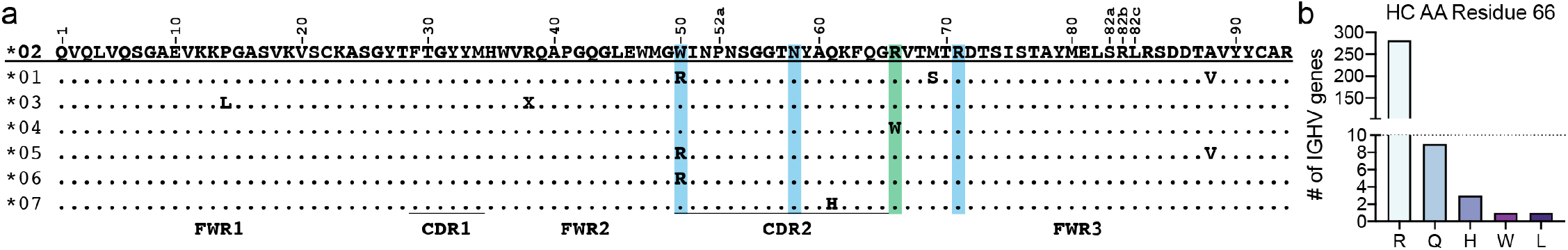
Allelic variations in IGHV1-2 genes impact CD4bs recognition and possibly BCR expression. **(a)** Amino acid sequence alignment of the seven IGHV1-2 alleles. The IGHV1-2*02 allele is shown as the reference. The Kabat numbering scheme of the residues is shown. Residues highlighted in blue indicate the position of key residues in IGHV1-2 required for binding to the CD4bs. Residue position 66 is shown in green. **(b)** Of all the IMGT curated functional IGHV genes (n=296), the majority of the genes utilized an arginine residue at position 66.

### VH1-2 allele frequency varies among human populations and is associated with variable usage in the naïve IgM repertoire

In light of these results, we sought to explore additional IGHV1 −2 allele signatures at the population level. To survey inferred allele and genotype frequencies at SNPs within IGHV1 −2 among different population subgroups, we first leveraged naïve B cell-derived transcriptomic data from the DICE cohort (database of immune cell expression, expression quantitative trait loci and epigenomics)^48^, representing an ethnically diverse collection of donors (n=75; African American, n=5; Asian, n=17; Caucasian, n=34; Hawaiian/Pacific Islander, n=3; Mixed ethnicity, n=15; Unknown, n=1). We focused our analysis on three primary single nucleotide polymorphisms (SNPs; rs1065059, rs112806369, and rs12588974) within this dataset, which differentiate IGHV1 −2 alleles *02, *04, *05, and *06 (Fig. 5a; Supplementary Fig. 3). Using individuals within this cohort with sufficient RNAseq reads mapping to these positions, we inferred individual SNP genotypes (Supplementary Fig. 3) and IGHV1 −2 allele-based genotypes (Fig. 5; Supplementary Table 3). Based on allele inferences, *02, *04, and *06 alleles were observed at frequencies of 42%, 43%, and 13% (Supplementary Table 1). The *02/*02 (18.6%), *02/*04 (34.6%), and *04/*04 (18.6%) genotypes were most common, followed by *02/*06 (12%) and *04/*06 (14.6%) heterozygotes (Fig. 5a). Evidence for the *05 allele was limited (1/75 individuals), and no *05/*05 homozygotes were observed in this cohort (Fig. 5a). Similarly, we did not observe SNP alleles representing *01, *03, or *07 (Supplementary Fig. 3). It was notable that in contrast to *05, both the *06 and *04 alleles were more prevalent in the population (Fig. 5a). Specifically, individuals lacking the *02 allele, represented by *04/*04 and *04/*06 genotypes, were observed at a collective frequency of ~33% in the overall cohort. Frequencies of these genotypes were moderately higher in both Caucasian (41%) and Mixed (40%) subgroups (Fig. 5a). In our eOD-GT8^+^ BCR sequencing data, we observed that the *04 allele was associated with significantly lower VRC01 -class B cell precursor frequencies relative to *02 among presumed *02/*04 heterozygotes (Fig. 1 and 2, Tables 1 and 2). Therefore, the VRC01-class B cell priming efficacy in *04 individuals may be reduced relative to *02 allele harboring individuals. Importantly, the *06 allele is represented by an arginine at AA position 50 which may hamper its potential to become a VRC01 -class bnAb as in the *05 allele.

**Figure 5.**
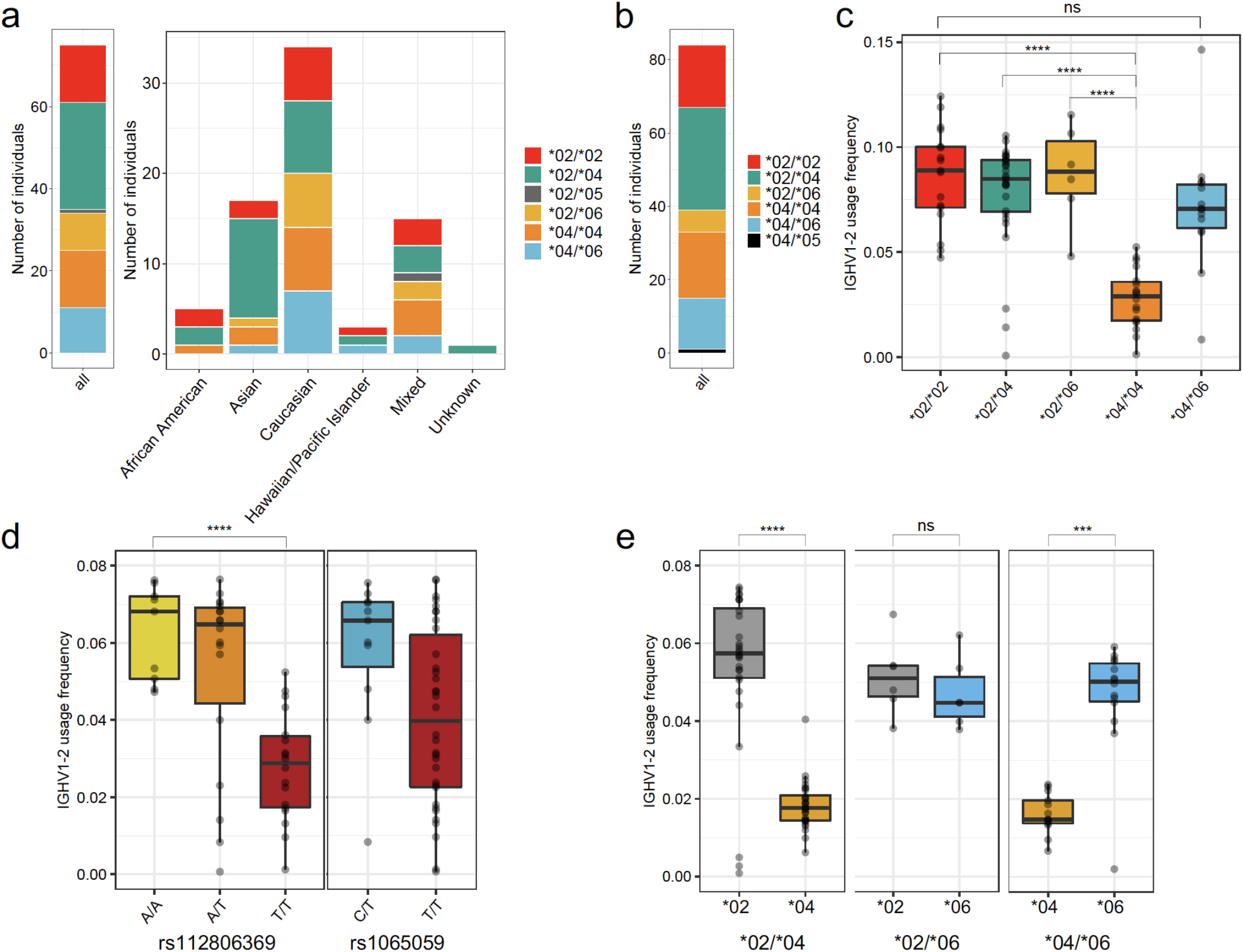
Population-level IGHV1-2 genotype frequencies and allele-specific effects on naïve antibody repertoire usage. **(a)** Number of individuals of each inferred IGHV1 −2 genotype in the collective cohort (left), and within each population subgroup (right), based on allele inferences made using RNA-seq data (n=75 donors). **(b)** Number of individuals of each IGHV1-2 genotype based on inferences made from IgM/IgD expressed antibody repertoire sequencing data (n=84 donors). **(c)** IGHV1-2 gene usage within the IgM/IgD repertoire partitioned by IGHV1-2 genotype (n=83 donors). **(d)** IGHV1 −2 gene usage in the IgM/IgD repertoire partitioned by genotypes at SNPs rs112806369 and rs1065059 (n=83 donors). **(e)** IGHV1-2 allele-specific usage within the IgM/IgD repertoire in IGHV1-2 heterozygotes (*02/*04, n=28 donors; *02/*06, n=6; *04/*06, n=14). One-way ANOVA: ***P<0.0001; ****P<0.00001; ns = not significant.

We additionally assessed allele frequencies at each these three SNPs in data from the 1000 Genomes Project (1 KGP^49^), which had been done previously when less data was available^13^. While technical confounding factors related to the use of short-read mapping and cell-line artifacts are known to influence the accuracy of genotype frequencies in the 1 KGP dataset^25,50^ (Rodriguez et al. *in prep),* requiring that these data be interpreted with caution, IGHV1 −2 SNP allele frequency biases are observable among human subpopulations (Supplementary Fig. 4). For example, consistent with the cohort studied here, the 1 KGP dataset also provides evidence that minor alleles at two SNPs associated with non-*02 alleles (rs112806369, *04; rs1065059, *05/*06) are relatively common across populations (14.9-46.4%). In comparison, while the SNP allele representing valine at position 86, observed in alleles *01 and *05 (rs12588974), occurs at lower frequencies in most populations (2.6-11%), it appears to be more common in the East (38.6%) and South Asian (22.6%) subpopulations (Supplementary Fig. 4). The fact that this contrasts with the limited support for *05 in the RNAseq dataset analyzed here could be explained by the smaller population subgroup sizes, as well as known expression biases in *05^51^ that may make it more difficult to detect from RNAseq data. These observations warrant more comprehensive sequencing of IGHV1 −2 as a means to fully clarify the extent of population-level germline variation at this locus.

Previous reports have identified allele-associated IGHV gene usage differences within naïve and antigen-stimulated B cell repertoires^51^”^55^. Given this, and the apparent preferential selection of the IGHV1 −2*02 allele observed in our VDJ sequencing data, we next investigated whether skewed IGHV1-2 allele usage patterns were observable within the naïve repertoire at the population level. To do this, we utilized a separate publicly available IgM/IgD expressed repertoire sequencing (RepSeq) dataset from a cohort of healthy donors (n=84)^56,57^. Consistent with genotype data reported above, among the individuals included in the RepSeq dataset, we observed the presence of IGHV1 −2*02, *04, and *06 alleles at the highest frequencies (Supplementary Table 3), represented by *02/*02 (20.2%), *04/*04 (21.4%), *02/*04 (33.3%), *02/*06 (7.1%), and *04/*06 (16.6%) genotypes (Fig. 5b). Only one heterozygous subject carried the *05 allele (*04/*05). The apparent low prevalence of *05 may occur due to low usage profiles of this allele^51^; the usage frequency observed in the *04/*05 individual was 1.3%. We observed a range in IGHV1 −2 usage frequencies across the remaining subjects (0.06-14%), which we found to associate with allelic variation (one-way ANOVA, effect of genotype, P = 3.76e-10; Fig. 5c). Consistent with observations discussed in above sections, *04/*04 homozygotes had the lowest IGHV1 −2 usage relative to all other genotypes (one-way ANOVA, P = 2.14e-10, vs. *02/*02; P = 2.82e-09, vs. *02/*04; P = 1.86e-07, vs. *02/*06; Fig. 5c). Usage in the *04/*06 heterozygotes, however, was not significantly different from *02/*02 (one-way ANOVA, P = 0.12). This effect of *04 on IGHV1-2 usage was clearly observable when we grouped subjects by genotype at SNP rs112806369 (Fig. 5d), which differentiates *04 (T) from *06 and *02 (A) alleles. Individuals of the A/A and A/T genotypes had higher IGHV1 −2 usage than those of the T/T genotype (one-way ANOVA, P = 1.46e-11; Fig. 5d). In contrast, partitioning subjects by genotypes at SNP rs1065059, which differentiates *06 (C) from *04 and *02 (T) did not reveal significant differences (one-way ANOVA, P = 0.21; Fig. 5d). Consistent with earlier observations^54,55^, we also observed *04 allele-specific usage biases within IGHV1 −2 heterozygotes (Fig. 5e). In individuals of *02/*04 and *04/*06 genotypes, *04 usage was significantly lower than that of the *02 (one-way ANOVA, P = 6.65e-12) and *06 alleles (one-way ANOVA, P = 0.0003). This was in contrast to allele-specific patterns in *02/*06 heterozygotes, in which both alleles were used at comparable frequencies (one-way ANOVA; P = 0.47). These results implied that the *02 usage bias relative to other IGHV1-2 alleles among VRC01-class bnAbs and naïve B cells likely occurs due to genetic impacts on V(D)J recombination frequencies and/or BCR expression.

## DISCUSSION

Previously we showed that naïve precursor B cells to different antigens can be identified by using fluorescent GT probes such as eOD-GT8, coupled with single cell Sanger sequencing^20,32^. Using the same eOD-GT8 probes but using a bulk-sort based, high-throughput single cell sequencing technology, we have confirmed that the human B cell repertoire can be quickly screened for rare antigen-specific naive B cells. The VRC01-class naïve B cell frequencies calculated based on sequences derived from the 10x Genomics Chromium platform were comparable to our previous numbers determined by Sanger sequencing. In this study, we used two different probes: SA tetramers and 60mer nanoparticles. Regardless of the probe used, the final calculated precursor frequencies of naïve VRC01 -class B cells were similar.

The eOD-GT8 60mer probes were found to be less efficient at enriching for VRC01-class naïve B cells than tetramer probes. The majority of 60mer sorted IGHV1 −2^+^ B cells were not paired with LCs with 5-AA LCDR3s. Compared to the tetramer, the 60mer probe was also less selective for IGHV1-2 overall. The affinity of the 60mer-isolated non-VRC01 -class naïve B cells here are unknown. Some of these BCRs identified by the 60mer probe, particularly those that are IGHV1 −2^+^, may have low monovalent affinity to the CD4bs that was enhanced by avidity. For example, we previously showed that non-VRC01 class IGHV1 −2 BCRs, and non-IGHV1 −2 BCRs expressed from naïve B cells isolated by tetramer probes had an average K_D_ of ~20 μM to eOD-GT8, which is in the range of low to medium affinity VRC01-class precursors.^32^ The increased proportion of non-IGHV1-2 BCRs could also be due to elevated representation of B cells that recognize regions outside of the CD4bs, but not picked up by the eOD-GT8^KO^ 60mer probe. If the latter case is true, specificity in sorting VRC01-class B cells by 60mer probes may be improved by adding the eOD-GT8^KO^ 60mer probe first. The same condition would also likely apply when using tetramer probes. Hence, our results here imply using tetramer probes coupled with 10x Genomics Chromium technology would be the most effective way to examine the B cell repertoire, although high avidity nanoparticle probes hold the potential for detecting low affinity B cells if strategies to further eliminate off-target B cells can be improved, and is worthy of further exploration.

In our analysis of eOD-GT8 probe binding VRC01 -class BCR sequences, we also made the observation that not all IGHV1 −2 germline alleles appear to make equal contributions to the circulating VRC01 -class B cell precursor pool, consistent with previously published work^36^. Specifically, we showed that the presence of the IGHV1 −2*02 allele within an individual’s genotype was associated with higher numbers of VRC01-class B cells. This was particularly true when comparing individuals harboring an *02 allele, compared to those with *04/*04 and *05/*05 genotypes. A nearly 2-fold reduction in VRC01 -class precursors was observed in the *04/*04 donor relative to those with an *02 allele. A complete absence of eOD-GT8 binding IGHV1-2 BCRs was observed in the *05/*05 donor. Further, in heterozygous *02/*04 individuals, representing 4/6 VRC01 -class naïve B cell positive donors, we observed that eOD-GT8-binding B cells were overwhelmingly associated with use of IGHV1 −2*02-derived BCRs. While the inability of *05 to contribute to VRC01 -class antibodies has been noted previously, it remains unclear whether the reduced precursor frequencies resulting from the *04 allele would impact the outcome of VRC01-class B cell priming immunizations. We noted that among all curated IGH alleles in IMGT, IGHV1 −2*04 is one of the few IGH alleles not encoding an arginine at AA position 66, and the only allele encoding a tryptophan at this position. It is plausible that W66 has potential functional consequences for BCR expression or solubility.

Interestingly, mirroring observations from our single-cell BCR analyses, we also noted strong genetically-driven usage biases of IGHV1 −2 alleles in the naïve repertoire of an expanded cohort of healthy donors. These analyses showed that, while *02 usage was relatively high in the overall naive repertoire, the *04 allele was utilized at very low frequencies in both homozygous and heterozygous individuals. In addition, although no donors in our eOD-GT8 probe sorting studies carried the *06 allele, we did find that the *06 allele was utilized at relatively high frequency within the naïve repertoire in our bulk RepSeq analysis, at levels comparable to *02. Because this allele lacks the critical W50 residue present in *02, it is predicted to not contribute to VRC01 -class antibodies. Whether being heterozygous for *02/*06 or *04/*06 would impact the frequency of VRC01 -class B cell precursor pool should be directly tested in future studies.

Together, these functional data indicated that inter-individual variation in GT vaccine responses, driven by differences in IGHV1 −2 genotype, could be expected. With this in mind, we also investigated the frequencies of IGHV1 −2 alleles and genotypes at the population level. Principally, this analysis revealed that both *04 and *06 alleles are frequent, and individuals completely lacking IGHV1-2*02 in their genomes make up a significant fraction of the population. In particular, the distribution of IGHV1 −2 alleles stratified by ethnic groups revealed differences that should likely be considered when developing vaccines.

In summary, we emphasize that a primary consideration in developing vaccines should be whether B cells that are to be targeted by immunogens exist within the naïve B cell repertoire of most people in a population of interest, and if those B cells occur at a high enough precursor frequency such that they are likely to become activated in response to immunizations. Better understanding of the factors that contribute to variation in naïve B cell precursor frequencies and repertoires will be critical moving forward. As illustrated in this study, the antigen-specific naïve B cell repertoire can be examined relatively quickly with state-of-the-art sequencing technologies, and can be incorporated to iterative immunogen design pipelines to advance vaccine discovery.

## MATERIALS AND METHODS

### Probe preparation

Avi-tagged eOD-GT8 and eOD-GT8^KO^ monomers, and eOD-GT8 and eOD-GT8^KO^ 60mer nanoparticles were recombinantly expressed in HEK293F cells by transient transfection and purified as summarized elsewhere^13^. The eOD-GT8^KO^ probes are eOD-GT8^KO II^ probes described in our previous study^32^, renamed for simplicity. Avi-tagged monomer probes were biotinylated and purified as previously described^29^. To generate eOD-GT8 tetramer probes, biotinylated monomers were mixed with fluorescently labeled streptavidin (SA-Alexafluor647 or SA-Brilliant Violet 421) at a molar ratio of 4 monomers: 1 SA, in a stepwise manner. 1/3 of the total amount of SA to be added to the biotinylated probes and incubated for 20 minutes in the dark at RT, and the process was repeated twice. The knock-out probe, eOD-GT8KO:SA-phycoerythrin (PE) was prepared in the same manner. eOD-GT8 60mer nanoparticles were directly labeled with fluorophores using AlexaFluor488 or Alexafluor647 protein labeling kits (Life Technologies) according to instructions supplied by the manufacturer. eOD-GT8KO 60mer nanoparticles were labeled with Pacific Blue protein labeling kit (Life Technologies).

### Sorting and 10X Genomics VDJ sequencing

Frozen PBMCs isolated from blood were thawed and recovered in R10 (RPMI, 5% FBS, 1x PenStrep, 1x Glutamax), and stained for sorting as previously described^32^. In brief, total PMBCs were enriched for B cells using a MACS human B cell isolation kit (Miltenyi Biotec). Purified B cells were enumerated, and stained for 20 min at 4 °C with a mix of tetramer or 60mer probes (two eOD-GT8 probes and one eOD-GT8^KO^ probe) in R10. Without washing, antibody master mix was added to the cells for an additional 20 min at 4 °C. Cells were washed twice and passed through a 70 μm mesh filter prior to sorting.

Where indicated, some total PMBCs were stained with Alexafluor647-eOD GT8 60mer for 20 min at 4 °C. Cells were washed twice, then positively selected for AlexaFluor647^+^ cells using Anti-Cy5/Anti-AlexaFluor647 MicroBeads (Miltenyi Biotec). The AlexaFluor647-enriched cells were stained for 20 min at 4 °C with AlexaFluor488-eOD GT8 60mer and Pacific Blue-eOD-GT8^KO^ 60mer. Without washing, antibody master mix was added for 20 min at 4 °C. Cells were washed twice and passed through a 70 μm mesh filter prior to sorting.

In some cases, TotalSeq-C Hashtag antibodies (Biolegend) were used to multiplex samples from different donors. 0.1 μg of Hashtag antibody per 1 million cells was added separately to each sample tube along with the antibody master mix.

All samples were sorted on a BD FACSAria II sorter using an 85 μm nozzle. eOD-GT8^+^ cells were gated as Lymphocytes/singlets/dump (anti-CD14, CD16, CD4, CD8, IgG, Live/Dead)/CD19^+^ or CD20^+^/eOD-GT8+eOD-GT8^+^/eOD-GT8^+^eOD-GT8^KOneg^, and bulk sorted into a 1.6 mL Eppendorf tube containing 50 μL of R10 catch buffer. In some cases, dump antibodies and Live/Dead were put on separate colors.

Sorted cells were spun down in a microcentrifuge, and extra buffer was removed until only approximately 30 μL was remaining in the tube. The pelleted cells were resuspended in 30 μL, and prepared following instructions provided for Chromium Single Cell V(D)J Reagent Kits with Feature Barcoding technology (10x Genomics). The legacy system was used for all experiments performed in this manuscript. VDJ cDNA libraries were sequenced on an Illumina MiSeq or NovaSeq 6000 using a 150×150 bp configuration, aiming for ~5000 read pairs per cell. Where Hashtag antibodies were used, Hashtag cDNA libraries were sequenced on the NovaSeq 6000 using the same configuration as for the VDJ library. Target number of hashtag reads was ~1600 read pairs per cell, amounting to approximately 1:3 Hashtag: VDJ library pooling ratio. Often, the actual number of cells recovered post cDNA generation was much fewer than the input number of cells estimated from the sort report, on average resulting in much higher number of read pairs per cell for the two libraries.

### BCR sequence analysis

The sequenced VDJ contigs were assembled and annotated using CellRanger VDJ within the CellRanger software package v3.1 (10X Genomics), using a VDJ reference library compiled from IMGT references. Each given cell barcode was associated with its productive heavy and light chain information. First, cells with only HC or LC contigs were removed from the dataset. Next, cell barcodes associated with multiple HC contigs were eliminated as this indicated that more than one cell was captured within a droplet. Barcodes with more than one LC contig of the same isotype were removed for the same reason. For cell barcodes that expressed one HC contig with one IGKV and one IGLV contig, it was assumed that the HC would be paired with the IGLV LC, because IGLV rearranges when IGKV cannot be coexpressed with the HC. In all of the samples, varying fraction of the cells were annotated as expressing class switched isotypes. All cells other than those annotated as expressing an IgM or IgD isotype HC were excluded from analysis.

Where relevant, Hashtag reads were enumerated using CellRanger count. Hashtag counts were associated with productive assembled VDJ sequences based on cell barcodes, and the information was outputted into a single file in tabular format. Hashtagged samples were manually deconvoluted based on the following Hashtag read count criteria; 1) The cell must have ≥1000 read pairs from a given Hashtag, and 2) <100 read pairs from all other Hashtags. For example, if a cell was associated with 5000 Hashtag 1 reads but also with 110 Hashtag 2 reads, the cell was considered to be contaminated and excluded from analysis. The Python script used to generate the tabulated data is available on GitHub (https://github.com/LJI-Bioinformatics/Filter-Cellranger-VDJ). Sequences of all VRC01-class precursor B cells are provided in Supplementary Tables 1 and 2.

### Sanger sequencing

The IGHV1 −2 locus was PCR amplified from genomic DNA (25 ng) of donor X using Qiagen HotStar HiFidelity Polymerase Kit (Catalog No. 202602), with previously published oligos (5’-GAGACTCTGTCACAAACAAACCA-3’; 5’-GTGTGTTCTCTTTCTCATCTTGGA-3’). Thermocycler conditions included an initial incubation at 95°C for 5 minutes, followed by 30 cycles of: 94°C for 15 seconds, 60°C for 1 minute, 72°C for 1 minute, and final extension at 72°C for 10 minutes. The resulting PCR product was cloned using the TOPO™ TA Cloning™ Kit, with One Shot™ TOP10 Chemically Competent E. coli (Catalog Number K4575J10). Briefly, TOPO cloning reactions were prepared for each PCR product using the manufacturer’s protocol. Five colonies were selected for Mini-Prep (Catalog No. 27104), and extracted DNA was Sanger sequenced using T7 and SP6 oligos. Allele sequences were confirmed by visual inspection of sequence chromatograms (Supplemental Fig. 2).

### Population-level genotype analysis

Naïve B cell RNA-seq were mapped to the hg19 reference genome using TopHap v1.4.1^58^ as part of a previously published study^48^. RNA-seq “.bam” files were obtained from this study, and the software package SAMtools^59^ was used to assess read depth and allele calls at SNPs representing each of the seven currently curated IGHV1-2 alleles (see Figure 4). To infer alleles and genotypes at each position, we required a total read depth (>3) and allele-specific read depth >1; only base calls with quality scores >32 (Phred 66) were considered. Based on these filter criteria, only 75 individuals from this cohort had sufficient read data available. Only positions representing the *02, *04, *05, and *06 alleles exhibited variation between individuals (rs1065059, rs112806369, and rs12588974; Supplemental Fig. 3). IGHV1-2 allele-based genotypes were inferred based on combined genotype calls made at each of these three SNPs. Phase 3 variant call summary data from the 1KGP^49^ was obtained from the Ensembl genome browser (https://uswest.ensembl.org/).

### Population-level analysis of expressed IgM/IgG antibody repertoire sequencing data

We analyzed previously published naïve antibody repertoire sequencing datasets from 84 healthy human donors, derived from PBMC RNA (SRA: SRP16 1 83 9)^56,57^. For each sample in this dataset, we selected reads representing IgM/IgD isotypes, assigned these sequences to the closest germline IGHV and IGHJ genes, and computed CDR3s using the DiversityAnalyzer tool^60^. Germline genotypes for each sample, inferred by TIgGER^61^, were downloaded from VDJbase^57^. We excluded individuals in which >2 IGHV1 −2 alleles were inferred. To minimize the impact of sequencing and amplification errors, we collapsed IgM/IgD sequences within an individual with identical CDR3s and computed usage the frequency of IGHV1-2 as the number of collapsed sequences aligned to IGHV1-2 within each individual normalized by the total number of collapsed sequences; allele-specific usage frequencies in individuals heterozygous for IGHV1-2 alleles were computed in the same way. Because we used only IgM/IgD sequences, we considered that clonal expansion would have little to no effect on the data. Statistical associations between IGHV1 −2 gene/allele usage and IGHV1-2/SNP genotypes were assessed using one-way ANOVA; the single *04/*05 individual was excluded from these analyses.

## ACKNOWLEDGEMENTS

We thank the LJI Flow core for sorting assistance, the LJI sequencing core for NovaSeq-6000 operation. We also thank D. R. Burton (The Scripps Research Institute) for allowing us to use the 10X Genomics Chromium Controller, and Y. Kato (LJI) for sharing some of the fluorescently labeled eOD-GT8 60mer probes. We thank K. Fung and J. Greenbaum (LJI) for providing scripts to compile Hashtag counts with VDJ annotated sequences. We also thank A. Madrigal, P. Vijayanand, and B. Schmiedel (LJI) for providing processed RNA sequencing data from the DICE cohort, used for our population-level analysis. This work was funded in part by grants from the National Institutes of Health, AI100663 (Scripps Center for HIV/AIDS Vaccine Immunology and Immunogen Discovery), AI144462 (Scripps Consortium for HIV/AIDS Vaccine Development) (S.C. and W.R.S.), R21AI142590 (C.T.W.), and R24AI138963 (C.T.W.); and by the International AIDS Vaccine Initiative (IAVI) Neutralizing Antibody Consortium (NAC) and Center (W.R.S.). The FACSAria II Cell Sorter was acquired through the NIH Shared Instrumentation Grant (SIG) Program (S10 RR027366).

## AUTHOR CONTRIBUTIONS

J.H.L., C.T.W., C.H.-D., S.C. designed the research. J.H.L., L.T., and J.T.K. performed experiments. J.H.L., J.T.K, Y.S., C.T.W., C.H.-D., S.C. analyzed data. W.R.S. provided immunogen probes. J.H.L., C.T.W., S.C. wrote the manuscript. All authors were asked to provide comments.

