## Supplementary for "Vaccine genetics of IGHV1-2 VRC01-class broadly neutralizing antibody precursor naïve human B cells"

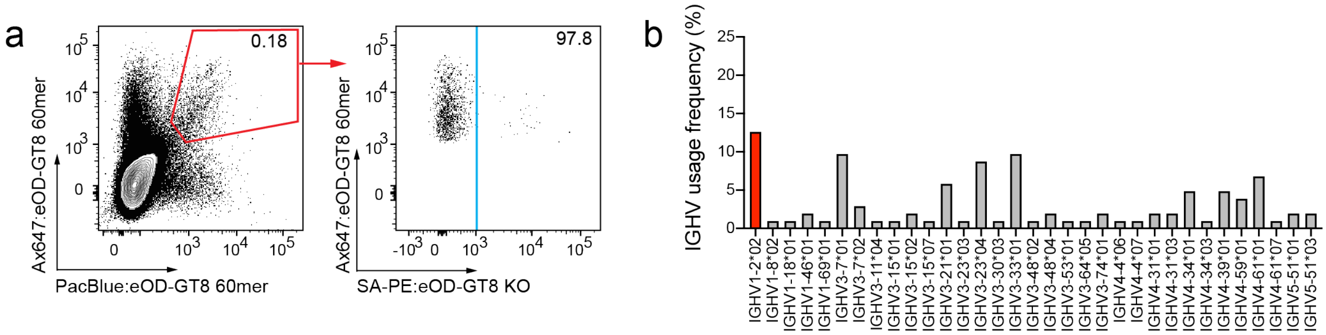


c

| LC | LCDR3 Length |
|---|---|
| IGKV1-39 | 10 |
| IGKV3-11 | 7 |
| IGKV3-15 | 10 |
| IGKV3-15 | 5 |
| IGKV4-1 | 9 |

**Supplementary Figure 1. Single cell sorting of eOD-GT8 60mer**

B cells enriched from healthy donor 7 PBMCs were stained with eOD-GT8 60mer probes directly conjugated with fluorophores and single cell sorted for eOD-GT8 60mer$^{+}$eOD-GT8$^{KOneg}$ naïve B cells.

**(a)** Flow cytometry of probe-staining cells.

**(b)** IGHV gene usage distribution among sequenced B cells (n=103).

**(c)** Successfully recovered LC sequences paired with BCRs expressing IGHV1-2 HC. Only one has a short 5-AA LCDR3.

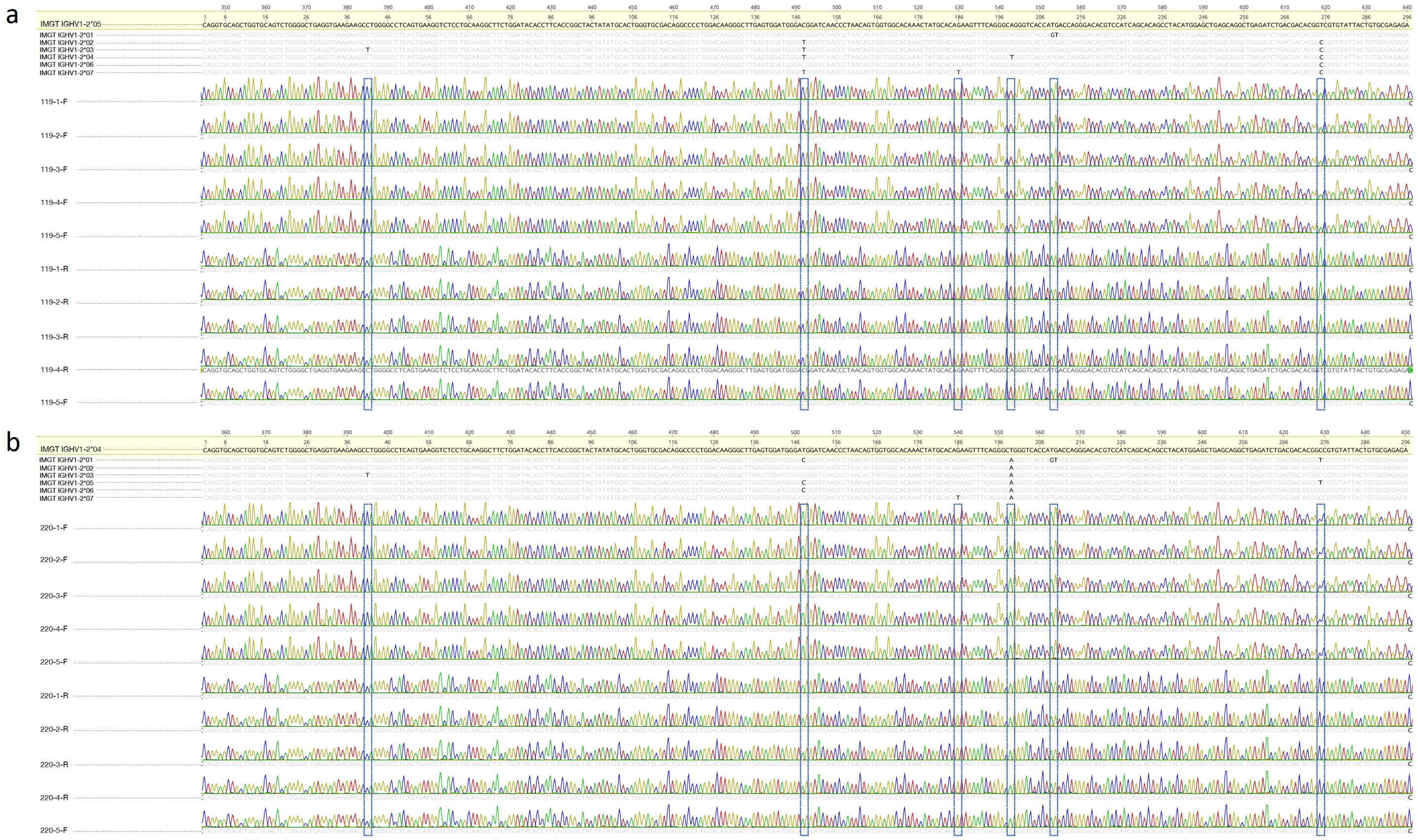

**Supplementary Figure 2. IGHV1-2 PCR, cloning, and Sanger sequencing confirm allele genotypes in donors 1 and 8**

Sanger chromatograms representing five sequenced clones from targeted IGHV1-2 gene PCR of genomic DNA in donors 8 **(a)** and 1 **(b)**. All clones were sequenced from both the forward and reverse directions to fully span the entirety of the IGHV1-2 coding region. Donor sequences are aligned to all known IGHV1-2 alleles, and SNPs differentiating either *05 or *04 from all other alleles are indicated (dashed boxes). As shown, all sequences within donor 8 **(a)** and donor 1 **(b)** matched *05 and *04, respectively, with 100% sequence identity.

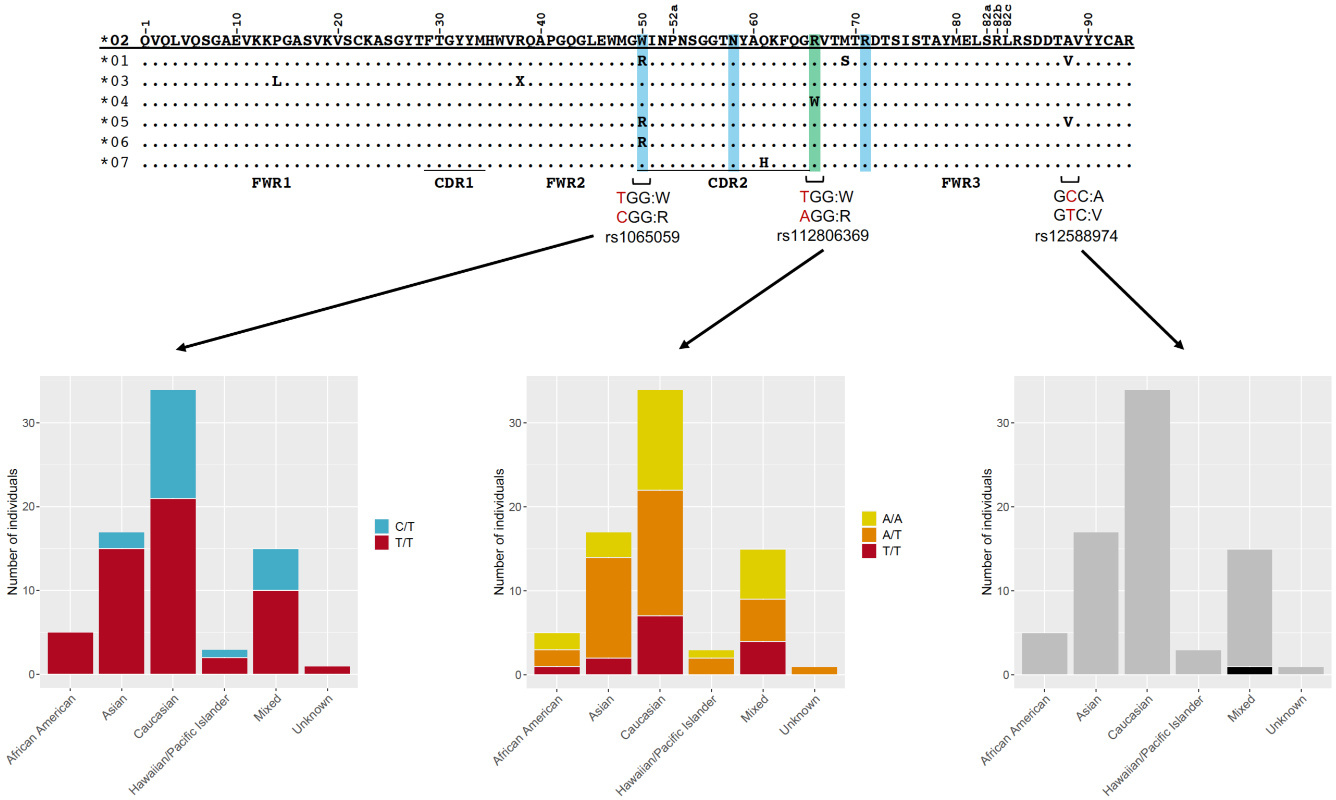


**Supplementary Figure 3. IGHV1-2 SNP genotypes inferred from RNA-seq data**

AA sequence alignment of the seven known IGHV1-2 alleles, as in Fig. 4a. AA positions represented by SNPs (rs1065059, rs112806369, and rs12588974) present in the RNA-seq data are indicated. Stacked bar plots showing the distribution of inferred SNP genotypes within each population subgroup (n=75; African American, n=5; Asian, n=17; Caucasian, n=34; Hawaiian/Pacific Islander, n=3; Mixed ethnicity, n=15; Unknown, n=1).

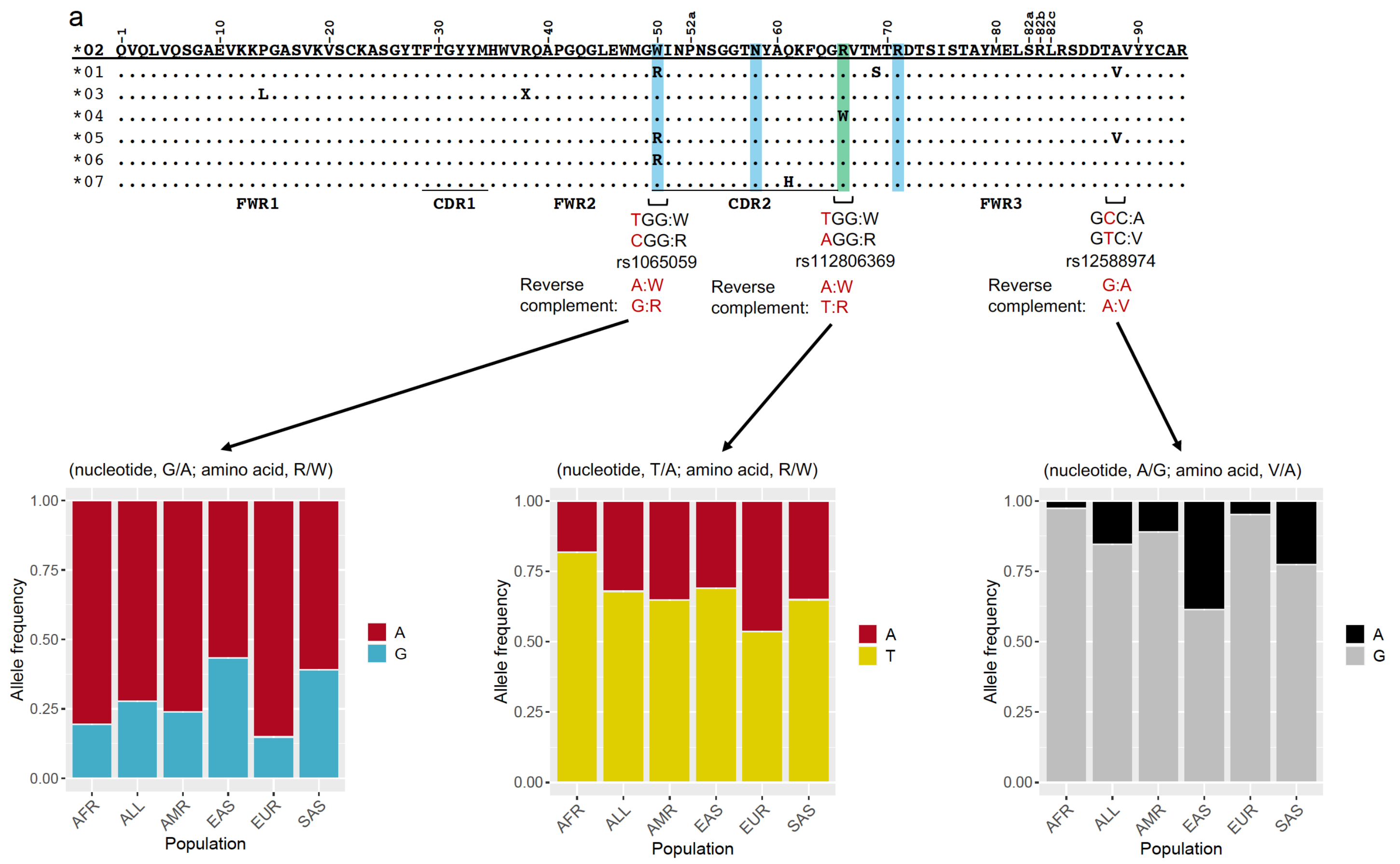


**Supplementary Figure 4. IGHV1-2 SNP allele frequency distributions in human subpopulations represented in 1000 Genomes Project data**

AA sequence alignment of the seven known IGHV1-2 alleles, as in Fig. 4a. AA positions represented by SNPs rs1065059, rs112806369, and rs12588974 are indicated. Reverse complement bases for each SNP are provided; the 1KGP variant call set contains complement alleles to reflect the fact that IGHV1-2 is found in reverse orientation (3'-5') in the genome reference assembly. Stacked bar charts displaying the allele frequencies at each SNP within the overall population, as well as five broad human subpopulations are shown (All, n=5,008; AFR, African, n=1,322; AMR, American, n=694; EAS, East Asian, n=1,008; EUR, European, n=1,006; SAS, South Asian, n=978).

| | IGHV | IGHD | IGHJ | CDRH3 | IGKV | IGKJ | CDRL3 |
|---|---|---|---|---|---|---|---|
| **D1-01** | IGHV1-2*04 | IGHD6-19*01 | IGHJ2*01 | CARVSIAVAGWYFDLW | IGKV3-20*01 | IGKJ1*01 | CQQYRTF |
| **D1-02** | IGHV1-2*04 | IGHD3-22*01 | IGHJ4*02 | CAREDDSSGYFYW | IGKV3-20*01 | IGKJ2*01 | CQQYQTF |
| **D1-03** | IGHV1-2*04 | IGHD4-17*01 | IGHJ2*01 | CARADYGDYWYFDLW | IGKV1-5*01 | IGKJ1*01 | CQQYETF |
| **D1-04** | IGHV1-2*04 | IGHD3-3*02 | IGHJ5*02 | CARATRGHFSFDPW | IGKV3-20*01 | IGKJ2*01 | CQQYETF |
| **D1-05** | IGHV1-2*04 | IGHD2-21*01 | IGHJ2*01 | CARDLGICGGDCYPYWYFDLW | IGKV3-7*01 | IGKJ1*01 | CQQYETF |
| **D1-06** | IGHV1-2*04 | IGHD6-6*01 | IGHJ3*02 | CARTSRLGDAFDIW | IGKV1-5*01 | IGKJ3*01 | CQQYETF |
| **D1-07** | IGHV1-2*04 | IGHD3-16*01 | IGHJ6*02 | CASGGGGGSGDYYYYGMDVW | IGKV3-20*01 | IGKJ2*01 | CQQYEAF |
| **D1-08** | IGHV1-2*04 | IGHD4-17*01 | IGHJ3*02 | CARDYGDPHAFDIW | IGKV3-20*01 | IGKJ2*01 | CQQYDTF |
| **D1-09** | IGHV1-2*04 | IGHD6-13*01 | IGHJ2*01 | CARDRGNSSSWHYWYFDLW | IGKV1D-33*01 | IGKJ4*01 | CQQYDSF |
| **D1-10** | IGHV1-2*04 | IGHD1-26*01 | IGHJ3*02 | CARGEYSGSYYAIW | IGKV1-5*01 | IGKJ1*01 | CQQPGTF |
| **D1-11** | IGHV1-2*04 | IGHD2-8*01 | IGHJ4*02 | CASGPKGLDYW | IGKV1D-33*01 | IGKJ3*01 | CQQFGAF |
| **D1-12** | IGHV1-2*04 | IGHD2-2*01 | IGHJ3*02 | CARVCSSTSCYGAFDIW | IGKV3-15*01 | IGKJ3*01 | CQQYNTF |
| **D1-13** | IGHV1-2*04 | IGHD6-19*01 | IGHJ4*02 | CARDRRLGSGWYFDYW | IGKV1D-33*01 | IGKJ3*01 | CQQYKTF |
| **D1-14** | IGHV1-2*04 | IGHD2-15*01 | IGHJ4*02 | CARGYEAAATYDYW | IGKV3-20*01 | IGKJ2*01 | CQQYGTF |
| **D1-15** | IGHV1-2*04 | IGHD1-26*01 | IGHJ3*02 | CARQHVSGSHAFDIW | IGKV3-20*01 | IGKJ1*01 | CQQYETF |
| **D1-16** | IGHV1-2*04 | IGHD4-17*01 | IGHJ4*02 | CARRSRDYGANVGYDYW | IGKV3-20*01 | IGKJ2*01 | CQQYEAF |
| **D1-17** | IGHV1-2*04 | IGHD3-22*01 | IGHJ3*02 | CAREYYYDRGKAGAFDIW | IGKV3-20*01 | IGKJ4*01 | CQQYDTF |
| **D1-18** | IGHV1-2*04 | IGHD3-22*01 | IGHJ5*02 | CARSRHSSASAFDPW | IGKV1D-33*01 | IGKJ2*01 | CQQYDTF |
| **D1-19** | IGHV1-2*04 | IGHD6-19*01 | IGHJ4*02 | CARSTEDSSGWYGYW | IGKV1-5*01 | IGKJ4*01 | CQHYGAF |
| **D1-20** | IGHV1-2*04 | IGHD1-26*01 | IGHJ4*02 | CARVGSYYDIYYFDYW | IGLV2-8*02 | IGLJ2*01 | CSYADLF |
| **D1-21** | IGHV1-2*04 | IGHD6-25*01 | IGHJ3*02 | CARFSSGGGGRNAFDIW | IGLV2-8*02 | IGLJ2*01 | CSYAALF |
| **D1-22** | IGHV1-2*04 | IGHD3-10*01 | IGHJ4*02 | CASRGSGSYYETW | IGLV2-13*02 | IGLJ1*01 | CSSSEVF |
| **D2-01** | IGHV1-2*04 | IGHD3-22*01 | IGHJ3*02 | CARAGNYDSSGYYRPAFDIW | IGKV1-39*01 | IGKJ4*01 | CQQSYTF |
| **D2-02** | IGHV1-2*04 | IGHD3-3*01 | IGHJ4*02 | CARASNYDFWSGYTYW | IGKV1-5*01 | IGKJ3*01 | CQHPETF |
| **D2-03** | IGHV1-2*04 | IGHD4-17*01 | IGHJ2*01 | CARGDYGDYLGHSDHAWYFDLW | IGKV3-20*01 | IGKJ2*03 | CQQYEGF |
| **D2-04** | IGHV1-2*04 | IGHD1-26*01 | IGHJ4*02 | CARGSGSYYDYW | IGKV1-9*01 | IGKJ4*01 | CQQLNSF |
| **D2-05** | IGHV1-2*02 | IGHD3-10*01 | IGHJ4*02 | CARMQYYGSGSYSGW | IGKV4-1*01 | IGKJ3*01 | CQQSETF |
| **D2-06** | IGHV1-2*02 | IGHD6-19*01 | IGHJ4*02 | CARVDSTRSSGWPIDYW | IGKV1-5*01 | IGKJ3*01 | CQQYNSF |
| **D2-07** | IGHV1-2*02 | IGHD5-12*01 | IGHJ4*02 | CARAVGSFRGYSGWKFDYW | IGKV3-20*01 | IGKJ5*01 | CQQYGSF |
| **D2-08** | IGHV1-2*02 | IGHD3-22*01 | IGHJ4*02 | CARVGFYYDSSGYYSYW | IGKV4-1*01 | IGKJ2*01 | CQQYYTF |
| **D2-09** | IGHV1-2*02 | IGHD3-10*01 | IGHJ5*02 | CARAVIEKLWFGELRGYWFDPW | IGKV1-5*01 | IGKJ3*01 | CQQYETF |
| **D2-10** | IGHV1-2*02 | IGHD6-13*01 | IGHJ3*02 | CAWQQLVNAFDIW | IGKV1D-33*01 | IGKJ3*01 | CQQYDFF |
| **D2-11** | IGHV1-2*02 | IGHD3-3*01 | IGHJ4*02 | CARGGVVIDYW | IGKV3-20*01 | IGKJ5*01 | CQQYGLF |
| **D2-12** | IGHV1-2*02 | IGHD5-12*01 | IGHJ4*02 | CARDKRPLVATIESGLYLDYW | IGKV1-5*01 | IGKJ4*01 | CQQYNSF |
| **D2-13** | IGHV1-2*02 | IGHD5-24*01 | IGHJ2*01 | CARVGGDGYKSYWYFDLW | IGKV3-20*01 | IGKJ1*01 | CQQYGTF |
| **D2-14** | IGHV1-2*02 | IGHD3-16*01 | IGHJ2*01 | CARLSKGGFGGGANWYFDLW | IGKV1D-33*01 | IGKJ4*01 | CQQYDSF |
| **D2-15** | IGHV1-2*02 | IGHD6-19*01 | IGHJ4*02 | CARIPLYSSGWYDYW | IGKV1-12*01 | IGKJ3*01 | CQQANSF |
| **D2-16** | IGHV1-2*02 | IGHD3-10*01 | IGHJ4*02 | CAREPGPLGEYGYW | IGKV1-9*01 | IGKJ5*01 | CQQLSTF |
| **D2-17** | IGHV1-2*02 | IGHD6-13*01 | IGHJ4*02 | CARDWGVDSSSWPGFDYW | IGKV3-20*01 | IGKJ1*01 | CQQYEAF |
| **D2-18** | IGHV1-2*02 | IGHD1-26*01 | IGHJ3*02 | CARWVGATTSDNDAFDIW | IGKV1-5*01 | IGKJ1*01 | CQQYNAF |
| **D2-19** | IGHV1-2*02 | IGHD6-19*01 | IGHJ4*02 | CARGRGRYFDWLFW | IGKV1D-33*01 | IGKJ4*01 | CQQYDNF |
| **D2-20** | IGHV1-2*02 | IGHD4-17*01 | IGHJ4*02 | CARYDYGDYWAFDYW | IGKV1-5*01 | IGKJ4*01 | CQQYRTF |
| **D3-01** | IGHV1-2*04 | IGHD2-15*01 | IGHJ4*02 | CARARDCSGGSCYEFSYW | IGKV1-5*01 | IGKJ1*01 | CQQYEAF |
| **D3-02** | IGHV1-2*04 | IGHD3-10*01 | IGHJ4*02 | CARGSTLWFGELLGKYFDYW | IGKV3-20*01 | IGKJ3*01 | CQQYDTF |
| **D3-03** | IGHV1-2*04 | IGHD1-26*01 | IGHJ2*01 | CARGGGGSYFTWYFDLW | IGKV3-20*01 | IGKJ4*01 | CQQYSTF |
| **D3-04** | IGHV1-2*04 | IGHD6-19*01 | IGHJ4*02 | CARYIAVAGTFDYW | IGKV1-5*01 | IGKJ2*01 | CQQFYTF |
| **D3-05** | IGHV1-2*02 | IGHD1-26*01 | IGHJ3*02 | CARGGSHGAFDIW | IGKV1-5*01 | IGKJ3*01 | CQQSSTF |
| **D3-06** | IGHV1-2*02 | IGHD6-13*01 | IGHJ5*02 | CARVTAAALFDPW | IGKV3-20*01 | IGKJ2*01 | CQHPYTF |
| **D3-07** | IGHV1-2*02 | IGHD6-6*01 | IGHJ4*02 | CASTQLAAWSFDYW | IGKV4-1*01 | IGKJ2*01 | CQLGNTF |
| **D3-08** | IGHV1-2*02 | IGHD6-6*01 | IGHJ2*01 | CARDRGGQLVGWYFDLW | IGKV1D-33*01 | IGKJ2*01 | CQQYFSF |
| **D3-09** | IGHV1-2*02 | IGHD3-22*01 | IGHJ3*02 | CAGNVYYYDSSGYYFWAFDIW | IGKV1-5*01 | IGKJ2*03 | CQQYNSF |
| **D3-10** | IGHV1-2*02 | IGHD3-22*01 | IGHJ3*02 | CARAPYDSSGYYYWAFDIW | IGKV3-20*01 | IGKJ2*01 | CQQFSFF |
| **D3-11** | IGHV1-2*02 | IGHD1-1*01 | IGHJ5*02 | CASNHRTYKQQLELSRWEFDPW | IGKV1-5*01 | IGKJ2*02 | CQQYGTF |
| **D3-12** | IGHV1-2*02 | IGHD2-15*01 | IGHJ4*02 | CARVGAAASWYFDYW | IGKV1D-33*01 | IGKJ4*01 | CQQYDNF |
| **D3-13** | IGHV1-2*02 | IGHD6-19*01 | IGHJ4*02 | CARYEQWLVWYYFDYW | IGKV3-15*01 | IGKJ2*01 | CQQYNNF |
| **D3-14** | IGHV1-2*02 | IGHD1-26*01 | IGHJ4*02 | CASRSNIVGAALGYW | IGKV3-20*01 | IGKJ4*01 | CQQYGSF |
| **D3-15** | IGHV1-2*02 | IGHD4-23*01 | IGHJ4*02 | CARFYGGNSCFDYW | IGKV1-12*01 | IGKJ4*01 | CQQANSF |
| **D3-16** | IGHV1-2*02 | IGHD3-22*01 | IGHJ4*02 | CARVLGNYYDSSGYYYGYW | IGKV3-20*01 | IGKJ1*01 | CQQFETF |
| **D3-17** | IGHV1-2*02 | IGHD1-26*01 | IGHJ2*01 | CARVLPLNGIVGATTGWYFDLW | IGKV3-20*01 | IGKJ2*02 | CQSLGTF |
| **D3-18** | IGHV1-2*02 | IGHD6-19*01 | IGHJ4*02 | CAREKGYSSGWSFDYW | IGKV4-1*01 | IGKJ4*01 | CQQYGTF |
| **D3-19** | IGHV1-2*02 | IGHD3-10*01 | IGHJ5*02 | CARARTGGSGSYVFDPW | IGKV1-5*01 | IGKJ2*01 | CQQYEFF |
| **D3-20** | IGHV1-2*02 | IGHD6-6*01 | IGHJ4*02 | CARDIIASSSSSESFWSFDYW | IGLV4-60*02 | IGLJ2*01 | CQTWALF |
| **D3-21** | IGHV1-2*04 | IGHD4-23*01 | IGHJ4*02 | CARDNPPFDFDYW | IGLV2-23*01 | IGLJ2*01 | CCSYVVF |
| **D3-22** | IGHV1-2*02 | IGHD5-12*01 | IGHJ4*02 | CARGDSGYDFDYW | IGLV2-14*02 | IGLJ3*02 | CSSYKVF |

**Supplementary Table 1. VRC01-class naïve precursor B cell sequences from tetramer donors**

| | IGHV | IGHD | IGHJ | CDRH3 | IGKV | IGKJ | CDRL3 |
|---|---|---|---|---|---|---|---|
| **D4-01** | IGHV1-2*02 | IGHD3-16*01 | IGHJ5*02 | CARPLTFGGVMMRNWFDPW | IGKV1-5*01 | IGKJ1*01 | CQQYTTF |
| **D4-02** | IGHV1-2*02 | IGHD3-3*01 | IGHJ2*01 | CARDPNVESDWYFDLW | IGKV1-5*01 | IGKJ1*01 | CQQYTTF |
| **D4-03** | IGHV1-2*02 | IGHD3-10*01 | IGHJ1*01 | CAREDGSGTEYFQHW | IGKV1-5*01 | IGKJ2*02 | CQQYSTF |
| **D4-04** | IGHV1-2*02 | IGHD6-13*01 | IGHJ4*02 | CARDLGIAAGFDYW | IGKV3-20*01 | IGKJ5*01 | CQQYNTF |
| **D4-05** | IGHV1-2*02 | IGHD2-2*01 | IGHJ6*02 | CARVSSLPYYYYYGMDVW | IGKV3-20*01 | IGKJ1*01 | CQQYGTF |
| **D4-06** | IGHV1-2*02 | IGHD3-22*01 | IGHJ4*02 | CARAGRYYDSSGYYQPFDYW | IGKV3-20*01 | IGKJ3*01 | CQQYGSF |
| **D4-07** | IGHV1-2*02 | IGHD1-26*01 | IGHJ4*02 | CARALFGGATGLGPYYFDYW | IGKV1-5*01 | IGKJ1*01 | CQQYETF |
| **D4-08** | IGHV1-2*02 | IGHD3-10*01 | IGHJ5*02 | CARGRLRSGGANGFDPW | IGKV3-20*01 | IGKJ1*01 | CQQYETF |
| **D4-09** | IGHV1-2*02 | IGHD1-7*01 | IGHJ6*03 | CAVSAKLELRRYYYYYMDVW | IGKV3-15*01 | IGKJ2*01 | CQQYETF |
| **D4-10** | IGHV1-2*02 | IGHD1-26*01 | IGHJ4*02 | CARLYSGSYYGGYYFDYW | IGKV3-20*01 | IGKJ2*01 | CQQYALF |
| **D4-11** | IGHV1-2*02 | IGHD5-24*01 | IGHJ5*02 | CATTDGYNYWWFDPW | IGKV3-20*01 | IGKJ2*01 | CQQWDTF |
| **D4-12** | IGHV1-2*02 | IGHD3/OR15-3a*01 | IGHJ3*02 | CARPQDWSRAFDIW | IGKV1-5*01 | IGKJ1*01 | CQQSWTF |
| **D4-13** | IGHV1-2*02 | IGHD3-22*01 | IGHJ4*02 | CARPAYYYDSSGYYDYW | IGKV1-5*01 | IGKJ1*01 | CQQSGAF |
| **D4-14** | IGHV1-2*02 | IGHD3-10*01 | IGHJ5*02 | CARDFRDGDTMVRGVIWTFDPW | IGKV1-9*01 | IGKJ2*01 | CQQLETF |
| **D4-15** | IGHV1-2*02 | IGHD6-6*01 | IGHJ4*02 | CARPHIAARYYFDYW | IGKV4-1*01 | IGKJ2*01 | CQQEYTF |
| **D4-16** | IGHV1-2*02 | IGHD5-18*01 | IGHJ6*02 | CARRGYTLHYYYYYGMDVW | IGKV3-11*01 | IGKJ3*01 | CQQADTF |
| **D4-17** | IGHV1-2*02 | IGHD3-22*01 | IGHJ4*02 | CARYWAARPDDSSGYYYIDYW | IGKV1-27*01 | IGKJ1*01 | CQNPGTF |
| **D4-18** | IGHV1-2*02 | IGHD4-17*01 | IGHJ5*02 | CAREVRDDYGANGGWFDPW | IGKV1-5*01 | IGKJ4*01 | CQHPGTF |
| **D4-19** | IGHV1-2*04 | IGHD5-12*01 | IGHJ6*02 | CARGGYSGYDENYYYYYGMDVW | IGKV1-5*01 | IGKJ1*01 | CQQYETF |
| **D4-20** | IGHV1-2*02 | IGHD1-26*01 | IGHJ4*02 | CARDPVGATTHFDYW | IGLV3-1*01 | IGLJ2*01 | CQAWEVF |
| **D5-01** | IGHV1-2*02 | IGHD6-13*01 | IGHJ6*02 | CARAVAGSSWYRDYYYYGMDVW | IGKV1-5*01 | IGKJ1*01 | CQHHGAF |
| **D5-02** | IGHV1-2*02 | IGHD4-17*01 | IGHJ3*02 | CARDSTAYDAFDIW | IGKV1-5*01 | IGKJ1*01 | CQHLETF |
| **D5-03** | IGHV1-2*02 | IGHD3-22*01 | IGHJ3*02 | CARGVPYYDSSGYYWSIW | IGKV3-15*01 | IGKJ1*01 | CQHQGTF |
| **D5-04** | IGHV1-2*02 | IGHD6-19*01 | IGHJ4*02 | CARESFLIAVAGTTFDYW | IGKV1-5*01 | IGKJ2*01 | CQHQGTF |
| **D5-05** | IGHV1-2*02 | IGHD3-10*01 | IGHJ6*02 | CARDRIITMVRGVISHYYYGMDVW | IGKV1-5*01 | IGKJ1*01 | CQQSGTF |
| **D5-06** | IGHV1-2*02 | IGHD6-19*01 | IGHJ4*02 | CARDFSGWYGYW | IGKV3-20*01 | IGKJ1*01 | CQQSGTF |
| **D5-07** | IGHV1-2*02 | IGHD6-19*01 | IGHJ4*02 | CARDQEGYSSGWYGTW | IGKV3-20*01 | IGKJ1*01 | CQQYGAF |
| **D6-01** | IGHV1-2*02 | IGHD5-12*01 | IGHJ2*01 | CARPTRRGYDYDSWWYFDLW | IGKV1-39*01 | IGKJ3*01 | CQQSWTF |
| **D6-02** | IGHV1-2*02 | IGHD3-22*01 | IGHJ4*02 | CARDPPYDSSGYYRWGFDYW | IGKV3-20*01 | IGKJ1*01 | CQQYGTF |
| **D6-03** | IGHV1-2*04 | IGHD2-15*01 | IGHJ3*02 | CARWTRNCSGGSCYLYRSGAFDIW | IGKV1-5*01 | IGKJ2*01 | CQQHHTF |
| **D6-04** | IGHV1-2*04 | IGHD4-17*01 | IGHJ2*01 | CARTTGSSYWYFDLW | IGKV3-20*01 | IGKJ1*01 | CQQYASF |
| **D6-05** | IGHV1-2*02 | IGHD4-11*01 | IGHJ2*01 | CARPIHDYSNYNPDWYFDLW | IGLV3-1*01 | IGLJ1*01 | CQARNVF |

**Supplementary Table 2. VRC01-class naïve precursor B cell sequences from 60mer donors**

| Cohort/Subgroup | N | IGHV1-2*02 | IGHV1-2*04 | IGHV1-2*05 | IGHV1-2*06 |
|---|---|---|---|---|---|
| **African American**[a] | 5 | 0.6 | 0.4 | 0 | 0 |
| **Asian**[a] | 17 | 0.47058824 | 0.47058824 | 0 | 0.05882353 |
| **Caucasian**[a] | 34 | 0.38235294 | 0.42647059 | 0 | 0.19117647 |
| **Hawaiian/Pacific Islander**[a] | 3 | 0.5 | 0.33333333 | 0 | 0.16666667 |
| **Mixed ethnicity**[a] | 15 | 0.4 | 0.43333333 | 0.03333333 | 0.13333333 |
| **Unknown**[a] | 1 | 0.5 | 0.5 | 0 | 0 |
| **ALL (RNAseq)**[a] | 75 | 0.42666667 | 0.43333333 | 0.00666667 | 0.13333333 |
| **ALL (RepSeq)** | 84 | 0.4047619 | 0.4702381 | 0.00595238 | 0.11904762 |

**Supplementary Table 3. Allele frequency of IGHV1-2 alleles inferred from RNAseq and RepSeq data**

[a]Subgroups from RNAseq data
